# SPECTRA: Generalist framework for reconstruction of high-quality metabolic networks at large-scale from multi-omics data

**DOI:** 10.64898/2026.04.02.716249

**Authors:** Pavan Kumar S, Subasree Sridhar, Noor Alsmadi, Radhakrishnan Mahadevan, Nirav Pravinbhai Bhatt

## Abstract

Metabolic network databases often suffer from inconsistent pathways with blocked reactions. Furthermore, current reconstruction methods lack scalable ways to integrate multi-omics data and cannot flexibly combine diverse modeling pipelines. We introduce SPECTRA, a unified platform that reconstructs metabolic networks across multiple biological scales. SPECTRA uses distinct optimization formulations and achieves a 56-fold speed improvement over existing algorithms for flux consistency based model reconstruction. We demonstrated SPECTRA across three applications. First, we gap-filled 7,302 microbial reconstructions from the AGORA2 database, reducing blocked reactions from an average of 482 to 169 per model. Second, we modeled synthetic gut microbiota at the community scale with superior performance in reaction inclusion. Third, we extracted models for 1,479 cancer cell lines, capturing more cancer hallmark reactions than current methods. This framework provides a scalable and flexible approach for modeling metabolic networks, and generates testable hypotheses for machine learning applications in biology.

## 1 Introduction

Constraint-based metabolic modeling (CBM) has become a key framework for translating rapidly accumulating genomic information into genome-scale representations of metabolism. Over the past two decades, CBMs have evolved from single-cell bacterial reconstructions to dynamic multi-tissue [1] and sex-specific human models containing over 80,000 reactions [2]. Yet, large metabolic network databases and derived genome-scale models often contain substantial fractions of blocked reactions and inconsistent pathways, which limit their predictive power and biological interpretability. At the same time, reconstruction methods designed to improve model consistency and quality remain computationally expensive, preventing their routine application to the rapidly expanding number of reconstructions now available and severely limiting their scalability to complex multicellular systems. Further, current methods have an inherent limitation around unblocking reactions with multiple reversible reactions, a ubiquitous feature of metabolic networks (refer to Supplementary texts S1.1, S1.2, S1.3 and S1.4). These challenges are compounded by the rigidity of reconstruction pipelines that cannot flexibly combine diverse model reconstruction strategies. We briefly review these bottlenecks next.

Omics data have enabled the automated reconstruction of metabolic models from universal reconstructions, a comprehensive network containing all candidate reactions with balanced stoichiometry. Automated reconstruction involves selecting reactions supported by omics evidence and network connectivity. For instance, Genome-scale models (GEMs) are built from genomic or metagenome-assembled genome (MAG) data using tools such as CarveMe [3], BIT [4], gapseq [5], and metaGEM [6], where database models serve as universal reconstructions. These GEMs can then be refined into context-specific models (CSMs) by incorporating transcriptomic, proteomic, and metabolomic evidence to capture condition-specific metabolism [7–18]. For more targeted analyses, top-down approaches can reduce large networks to minimal reactomes (MRs), the smallest subnetworks that preserve key biological functions [19–21]. While these top-down approaches have improved the efficiency of unicellular model construction, applying them to multicellular systems such as microbial communities, multi-tissue networks, or host–pathogen models remains challenging. Current efforts typically rely on bottom-up strategies, where models for individual members are first reconstructed and curated, then combined into a shared simulated environment to approximate multicellular behavior [1, 2, 22–29]. However, this paradigm cannot leverage collective information from multicellular-level datasets and often overlooks interspecies metabolic interactions. Recent tools such as scFBA [30] and METAflux [31] integrate multi-omics data at the community level but operate as flux estimators, not model constructors. They predict flux distributions on pre-existing community models rather than constructing new, structurally refined models from universal reconstructions. Our previously developed community gap-filling algorithm [32] constructs models from universal reconstructions but has been demonstrated only for communities of up to three microbes.

Model reconstruction methods use diverse objectives tailored to specific biological questions. Omics evidence can be integrated by enforcing core reaction inclusion, optimizing fluxes with omics-based weights, or constraining input–output fluxes from exo-metabolomics data. These strategies either minimize network size (reaction count or total flux), balance inclusion of positively supported reactions against exclusion of negatively supported ones, or maximize biomass while limiting non-essential fluxes. An overview of representative methods, their objectives, and the constraints used to incorporate multi-omics data is provided in Supplementary Table S4. The ability to combine different objectives and constraints allows modelers to tailor the extraction process to their experimental needs and biological hypotheses. Some methods ensure biomass precursor production under topological constraints [33], while others enforce stoichiometric consistency in core reactions [17]. However, no existing approaches simultaneously ensure topological consistency of core reactions and the production of biomass precursors and is scalable. Likewise, although mixed-integer linear programming (MILP)-based approaches exist for identifying minimal microbiomes [34], analogous scalable linear programming (LP)-based methods are lacking. Furthermore, while some approaches combine task-based gap-filling with trade-off–based objectives [15], none integrate task-based gap-filling with minimal network–based objectives.

To address these limitations, we introduce SPECTRA (Scalable Platform for Extracting Constraint-based Top-down Reconstructions and Analysis), a general, scalable framework for extracting metabolic models from multi-omics data for a wide range of modeling applications. SPECTRA accepts omics evidence as input in the form of core reactions, reaction weights, and box constraints, and integrates these variables into universal reconstructions via highly customizable constraint and objective choices. SPECTRA uniquely supports alternative solution generation through both LP and MILP formulations, enabling the extraction of functionally equivalent models for complex biological scenarios. The platform comprises of eight optimization formulations, enabling the reconstruction of metabolic models at scales ranging from single cells to multicellular systems and entire microbial communities (Fig. 1). We applied SPECTRA across three key cases: (i) gap-filling 7,302 AGORA2 microbial reconstructions, (ii) community-scale modeling of synthetic gut microbiota, and (iii) context-specific modeling of 1,479 cancer cell lines. Using enhanced universal reconstructions like Recon3D+, a manually curated human model with improved gene-protein-reaction associations - SPECTRA ensures biological consistency across these diverse applications while demonstrating superior computational efficiency and biological accuracy compared to existing methods. The modular architecture unifies diverse reconstruction strategies, providing unprecedented flexibility for systems biology research. SPECTRA is implemented twice, as a MATLAB toolbox built on the COBRA Toolbox [35] and as a Python package built on COBRApy [36], so that it can be adopted from either ecosystem; the two share the same optimization formulations and interfaces, and each is accompanied by documentation and worked tutorials covering every formulation and use case described here. The results reported in this work were obtained with the MATLAB implementation, because the methods SPECTRA is benchmarked against are distributed only for MATLAB and a like-for-like runtime comparison requires all of them to run in a single environment.

**Fig 1.**
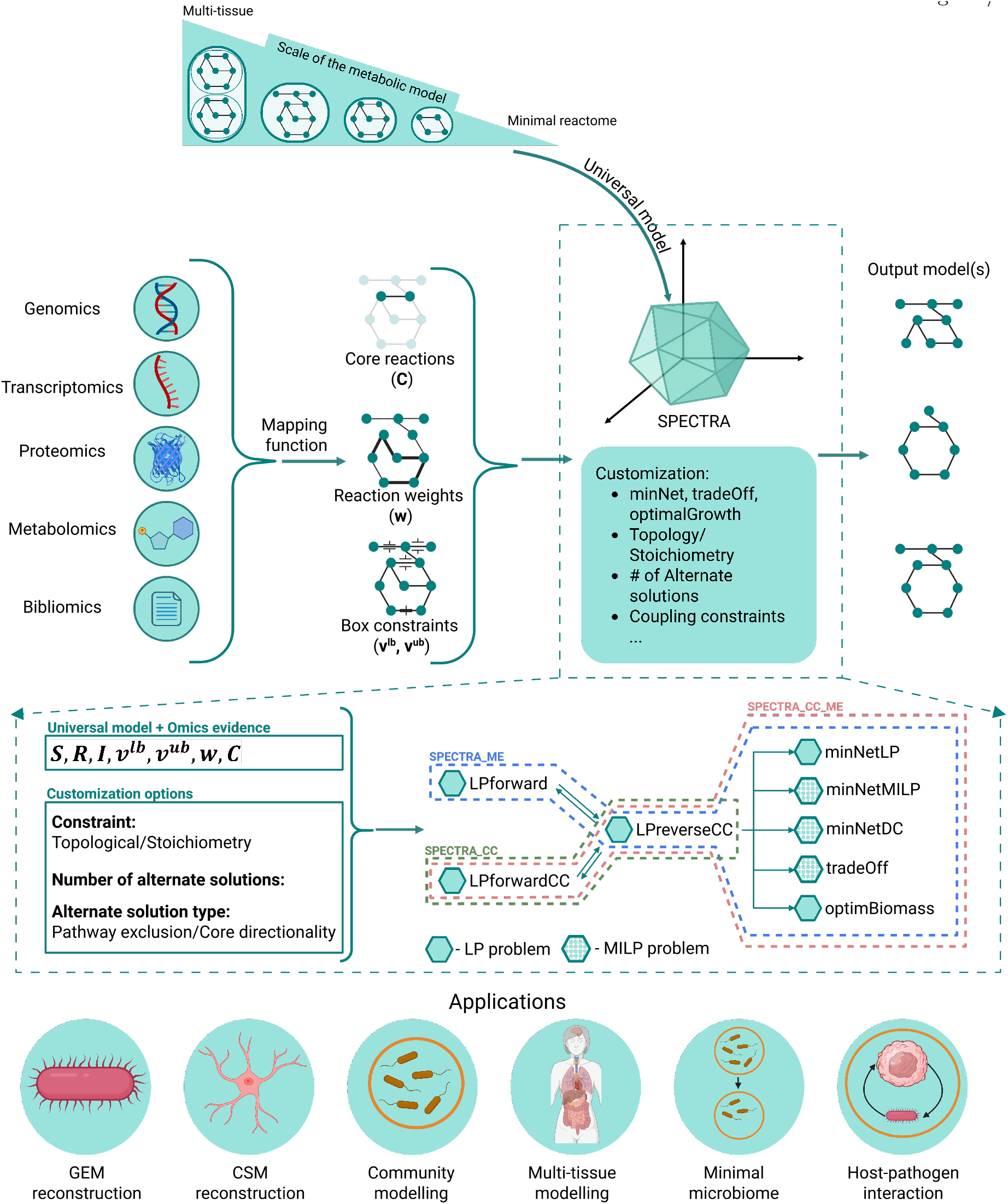
Overview of SPECTRA: SPECTRA is a unified, scalable framework that integrates omics data with universal reconstructions to extract metabolic models at multiple scales. The workflow begins by deriving core reactions (***C***), reaction weights (***w***), and box constraints (***v***^***lb***^ and ***v***^***ub***^) from omics data then customizing the constraints and optimization framework. SPECTRA comprises eight optimization formulations: three LP formulations (*LPforward, LPreverseCC, LPforwardsCC*) to identify a flux vector with non-zero flux through core reactions, and five network inference formulations that either minimize total flux (*minNetLP* ) or the number of reactions (*minNetMILP* ) or cardinality minimization using difference-of-convex functions (*minNetDC*), or balance inclusion of reactions with positive versus negative evidence (*tradeOff* ), or maximize biomass while limiting flux through other reactions (*optimBiomass*). SPECTRA finds its application in distinct model reconstructions that includes: GEM, CSM, Microbial community model building, Multi-tissue model building, Minimal microbiome inference and to study host pathogen interaction. (The figure is created using Biorender.com)

## 2 Results

### 2.1 Overview

SPECTRA provides a scalable, generic framework for integrating the multi-omics data (Fig. 1) to construct metabolic networks at various biological scales. The platform combines multiple optimization strategies (LP and MILP formulations) constrained by omics evidence on the universal reconstructions to optimize for metabolic network inference. By integrating diverse methods of optimization through a common framework, SPECTRA offers flexibility and control over the biological assumptions. This approach supports integrating multiple omics data to infer metabolic networks that range from minimal reactomes and single-cell context-specific models (CSMs) to complex multicellular systems.

All the metabolic network inference methods in SPECTRA follow a similar structure: loading a universal reconstruction, using a mapping function to derive core reactions, reaction weights, and box constraints from the omics data. These are used as optimization variables and parameters that determine which metabolic substructure is extracted. The derivation of the mapping function is outside the scope of this study and has been described elsewhere [3, 15, 18, 37, 38]. This unified framework reveals that many widely used metabolic network reconstruction methods are special cases of SPECTRA making it straightforward to extend or improve a wide range of methods.

The scalability of SPECTRA arises from an efficient strategy for identifying a flux vector with non-zero flux through core reactions that is consistent with the omics evidence. This is accomplished using three distinct LP formulations (described in detail in the Methods), after which the resulting flux vector is incorporated as additional constraints in the subsequent network inference step. This separation of a fast flux-finding phase from the more complex extraction formulations enables efficient performance even for thousands of models.

The generic nature of SPECTRA is rooted in the customization options it provides for network connectivity, the number and type of alternative solutions, and the choice of network inference formulation. Network connectivity can be enforced via stoichiometric constraints, assuming pseudo–steady state, or via purely topological constraints that consider only reaction connectivity. Multiple subnetworks can satisfy the same omics-constrained objective, and SPECTRA can systematically enumerate alternative optimal networks (and sub-optimal networks) using pathway exclusion (forcing at least one reaction to differ between solutions) or core reaction directionality (based on the sign of flux in reversible core reactions). SPECTRA currently implements five optimization formulations for network inference: minimizing total flux (*minNetLP* ), minimizing the number of reactions (*minNetMILP* ), cardinality-constrained minimization using difference-of-convex (DC) functions that approximates the *l*_0_-norm count of active reactions through a sequence of convex subproblems [39] (*minNetDC*), balancing inclusion of reactions with positive evidence against exclusion of reactions with negative evidence (*tradeOff* ), and maximizing biomass while minimizing flux through non-biomass reactions (*optimBiomass*). The specific utility and functionality of these formulations are shown in Supplementary text S1.5 where we apply the SPECTRA framework to a toy metabolic network. We next evaluate SPECTRA’s performance on minimal model generation and identifying blocked reactions in large metabolic models such as the human metabolic model Recon3D.

### 2.2 SPECTRA is computationally efficient for minimal model generation given a core reaction set

Model extraction algorithms such as Swiftcore, Fastcore, MBA, and mCADRE aim to retain all core reactions while minimizing the size of the universal reconstruction. Among these, MBA and mCADRE require performing a consistency check for every non-core reaction removal, which substantially increases their runtime relative to Fastcore and Swiftcore. To benchmark the computational efficiency of SPECTRA, we compared it against the LP-based methods Fastcore and Swiftcore. Depending on whether the universal reconstruction exhibits stoichiometric consistency, SPECTRA utilizes either the SPECTRA_ME framework (for consistent networks) or SPECTRA_CC_ME (for inconsistent networks). Initially we demonstrate efficiency of SPECTRA_ME framework with the *minNetLP* formulation for network inference. We used the human reconstruction Recon3D [40] as the universal reconstruction, and selected core reactions randomly in sizes ranging from 20 to 10,600. All three algorithms-SPECTRA_ME, Fastcore, and Swiftcore require a stoichiometrically consistent universal reconstruction as an input. Swiftcore has two variants: Swiftcore with network reduction (Swiftcore-R) and Swiftcore without reduction (Swiftcore-WR); here, we report results using Swiftcore-WR, while the results of Swiftcore-R are provided in Supplementary Fig. S9. As shown in Fig. 2a and S9, SPECTRA_ME solves fewer Linear Programs (LPs) than both Fastcore and Swiftcore. However, its average runtime (1.18 s) was slightly higher than that of Swiftcore (0.826 s). Upon analysis, we found that Swiftcore does not impose box constraints on non-core reversible reactions, which can result in infeasible models (see Supplementary Text S1.4). To test the effect of these constraints, we developed SPECTRA_ME2, a variant of SPECTRA_ME that only enforces irreversibility constraints. When tested on the same sets of core reactions, SPECTRA_ME2 achieved a lower average runtime (0.4864 s) than Swiftcore (Supplementary Fig. S10), thereby demonstrating the inherent efficiency of SPECTRA’s LP formulation.

**Fig 2.**
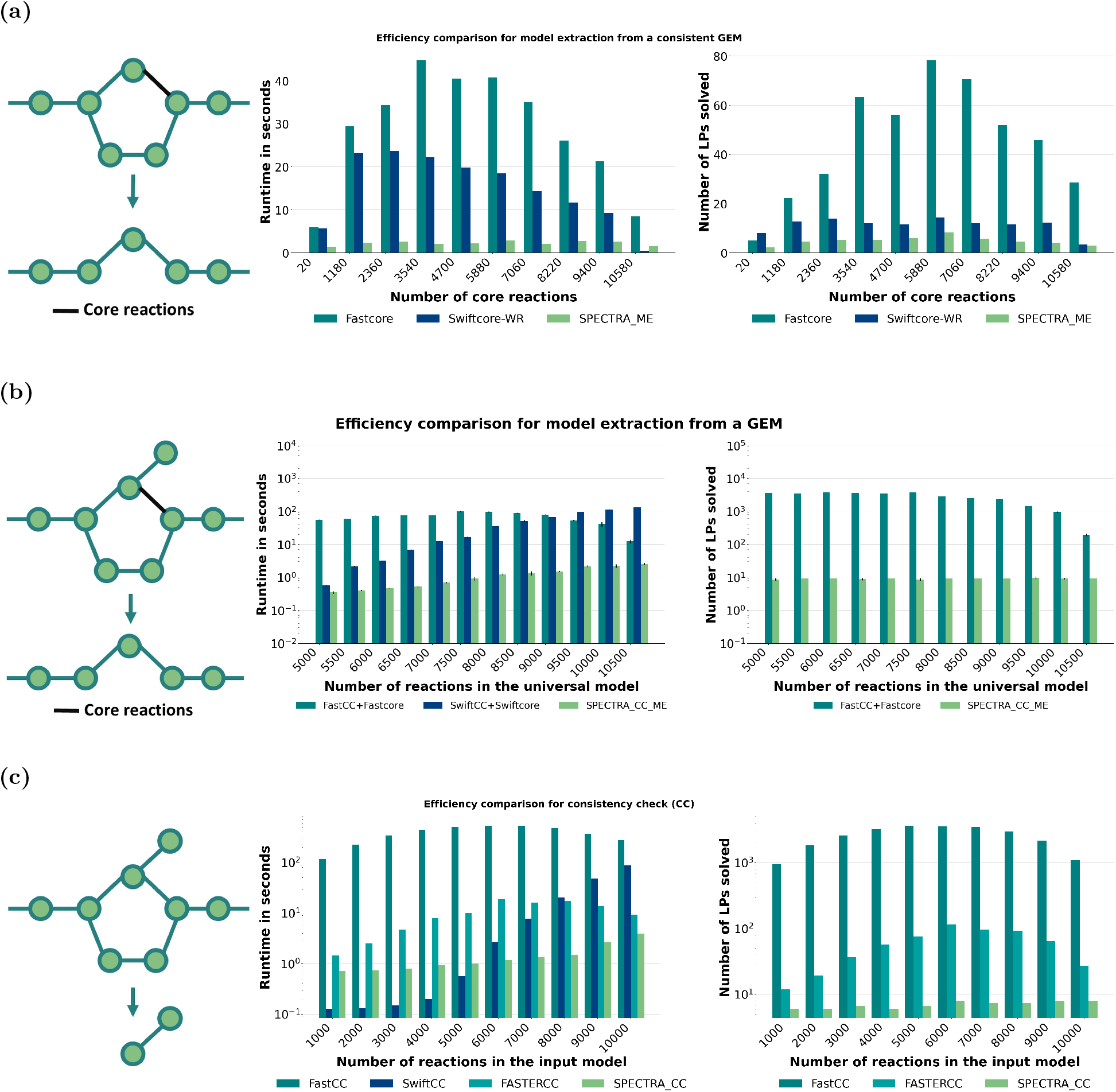
SPECTRA improves the efficiency of model extraction and blocked reaction identification. (2a) Runtime comparison of SPECTRA_ME, Fastcore, and Swiftcore on consistent universal reconstructions (Recon3D): SPECTRA_ME achieves the fastest extraction of minimal models across varying core reaction sizes. (2b) Performance of SPECTRA_CC_ME on inconsistent universal reconstructions: SPECTRA_CC_ME outperforms FastCC+Fastcore and SwiftCC+Swiftcore in both runtime and LP usage. Error bars indicate standard deviation of runtime across different core reactions. SwiftCC does not utilize LP to completely identify the blocked reactions. Hence comparison on number of LPs solved is done only with FastCC+Fastcore algorithm. (2c) Benchmarking of SPECTRA_CC against FastCC, SwiftCC and FASTERCC for blocked reaction detection on random subsets of Recon3D reactions. SPECTRA_CC solves the fewest LPs at every size. SwiftCC is faster on the smallest networks but scales poorly. SwiftCC is excluded from the LP comparison as it uses orthogonal–triangular decomposition to check the consistency of reversible reactions.

When the universal model is stoichiometrically inconsistent, Fastcore and Swiftcore require a separate consistency check to first remove inconsistent reactions. The standard extraction pipelines in such cases are FastCC+Fastcore and SwiftCC+Swiftcore, where consistency checking is performed first, followed by the removal of inconsistent core reactions, and finally model reconstruction. To address this limitation, we developed SPECTRA_CC_ME, which performs consistency checking and model extraction simultaneously (refer to Methods section). For benchmarking, we generated random universal models containing 5000–10,600 reactions, with randomly selected core sets of size 50–5000 in Recon3D. Because these universal sets are randomly constructed, they are inherently inconsistent. As shown in Fig. 2b, SPECTRA_CC_ME solves substantially fewer LPs than Fastcore. Moreover, runtime analysis shows that SPECTRA_CC_ME (average runtime = 1.19 s) is approximately 56 *×* faster than FastCC+Fastcore (66.68 s) and 36 *×* faster than SwiftCC+Swiftcore (43.92 s). Collectively, these results demonstrate that SPECTRA_ME and SPECTRA_CC_ME are highly efficient algorithms for reducing model size while ensuring the inclusion of all core reactions, and are especially advantageous when handling inconsistent universal reconstructions. Given this improved performance of SPECTRA on model extraction tasks, next we wanted to assess its performance on identification on blocked reactions in a constraint-based model.

### 2.3 SPECTRA is the fastest algorithm to get the blocked reactions

Blocked reactions are reactions in the model that cannot carry a non-zero flux under any feasible condition because of the structure or the constraints applied. Structural constraints can be of two types: stoichiometry or topology. Explanation of the different blocked reactions are provided in the Supplementary text S1.7. The existing algorithms deal with the stoichiometry structural constraints, and hence here we use the same for efficiency comparison. Several algorithms have been developed for the verification of stoichiometric consistency in metabolic models on a genome scale, including Flux Variability Analysis (FVA) [41], FastFVA [42], the CMC algorithm [7, 13], FastCC [7], SwiftCC [8], the cardinality minimization algorithm [39], and FASTERCC [43]. Among these, FastCC, SwiftCC and FASTERCC are reported to be computationally the most efficient [8,39,43], and we therefore benchmarked SPECTRA_CC against all three. Subsets of 1,000 to 10,000 reactions were drawn at random from the flux-consistent core of the human reconstruction Recon3D [40] and used as input networks. Each network size was generated as three independent random draws and each draw was timed three times; all four methods were run in a single environment, on one solver (Gurobi) at its default feasibility tolerance (10^*−*6^), so that the comparison is internally consistent (Methods 4.8).

SPECTRA_CC solved the fewest linear programs at every network size and was the only method whose LP count did not grow with the size of the network, requiring 6–8 LPs across the entire range against 12–116 for FASTERCC and 947–3,648 for FastCC (Fig. 2c). Summed over the ten network sizes, SPECTRA_CC was also the fastest method overall (14.8 s), ahead of FASTERCC (101.3 s, 6.8*×* ), SwiftCC (166.7 s, 11.3*×* ) and FastCC (3,824 s, 259*×*). The advantage is size-dependent rather than uniform. SwiftCC was faster on the smaller networks, up to 5,000 reactions, where the consistency check is dominated by a single matrix factorisation; but its runtime grew steeply with network size, and by 10,000 reactions it was 22*×* slower than SPECTRA_CC (87.6 s against 4.0 s). SPECTRA_CC was the fastest method at every size from 6,000 reactions upward, including against FASTERCC, whose best result was 9.2 s at 10,000 reactions.

Where the methods disagreed in the blocked reactions, we adjudicated by FVA. Across all instances only 12 reactions were classified differently by the four methods. SPECTRA_CC and SwiftCC each reported five of these as consistent when FVA proves them blocked, and FASTERCC reported two as blocked when FVA shows they carry flux. For SPECTRA_CC and FASTERCC this residual error is numerical: repeating the benchmark with the feasibility tolerance tightened to 10^*−*9^ removes it entirely, while SwiftCC continues to report blocked reactions as consistent (Supplementary Table S1).

SwiftCC identifies blocked irreversible reactions by solving a single LP but uses orthogonal–triangular decomposition for reversible reactions, so the comparison of the number of solved LPs was made only between SPECTRA_CC, FASTERCC and FastCC. One limitation of SwiftCC is that it only considers thermodynamic (irreversibility) constraints. It does not account for constraints on exchange reactions or reactions with positive lower bounds. However, CBMs often include additional constraints such as media composition, regulatory effects, ATP maintenance requirements, and enzyme capacity limitations. SPECTRA_CC and the other methods incorporates all such constraints, providing a more comprehensive and efficient approach for identifying blocked reactions.

The efficiency of SPECTRA_CC extends well beyond the scale shown here. On expanded Recon3D reconstructions, which represent multicellular systems as *k* replicate copies of the base network, it identified the consistent subnetwork of a 454,620-reaction model while solving only 10 LPs (Supplementary Text S1.9). At this scale the result also becomes sensitive to the solver tolerances: on microbial community models of up to 180,874 reactions, SPECTRA_CC returned a reproducible consistent set only at a feasibility tolerance of 10^*−*9^, repeated runs of the same model differing by up to 19,664 reactions at looser settings (Supplementary Text S1.10, Supplementary Table S2). On the whole-body reconstruction Harvey, whose bounds force 167 reactions to carry flux and so exclude the zero flux vector, the consistent subnetwork returned by a first pass is not itself a feasible model, and consistency checking must be iterated to converge (Supplementary Text S1.11, Supplementary Table S3).

Building on this foundation of rapid model building and consistency checking, we evaluated SPECTRA’s performance in reconstructing minimal, biologically meaningful metabolic networks from large universal models, assessing both computational performance and model quality.

### 2.4 SPECTRA derived models of microbes in AGORA database have fewer blocked reactions

The AGORA2 collection comprises of 7,302 genome-scale metabolic reconstructions representing more than 4,000 unique human intestinal microbial species [44]. Despite its comprehensiveness, stoichiometric consistency tests reveal that, on average, about 482 reactions are blocked in each model. We applied SPECTRA first to identify the most effective minimal-network–based inference method, and then used this method to perform gap filling on the AGORA2 models. Here, gap filling is entirely based on the stoichiometric structure of the universal reconstruction, with equal weights assigned to all non-core reactions.

In the first step, a random 30% of reactions were removed from each consistent AGORA2 model. These “reaction-deletion” models were then gap filled using a universal reconstruction constructed as the union of all reactions, metabolites, and their attributes (refer to Methods 4.9). The universal reconstruction was checked for conflicts; in particular, the reactions CINNMRi and COF420 NADP OX had conflict in reversibility constraints across models, and the reversible version was retained in the universal model. The gap-filled models were compared against the original consistent AGORA2 models. Performance metrics, including accuracy, precision, recall, and F1-score, are reported in Fig. 3a and Supplementary Fig. S11, with formulas provided in Methods 4.9. Overall, *minNetMILP* outperforms the other two methods in both accuracy and F1-score. While all methods achieve similar recall, *minNetMILP* shows higher precision, highlighting its superior ability to recover true positive reactions while minimizing false positives.

**Fig 3.**
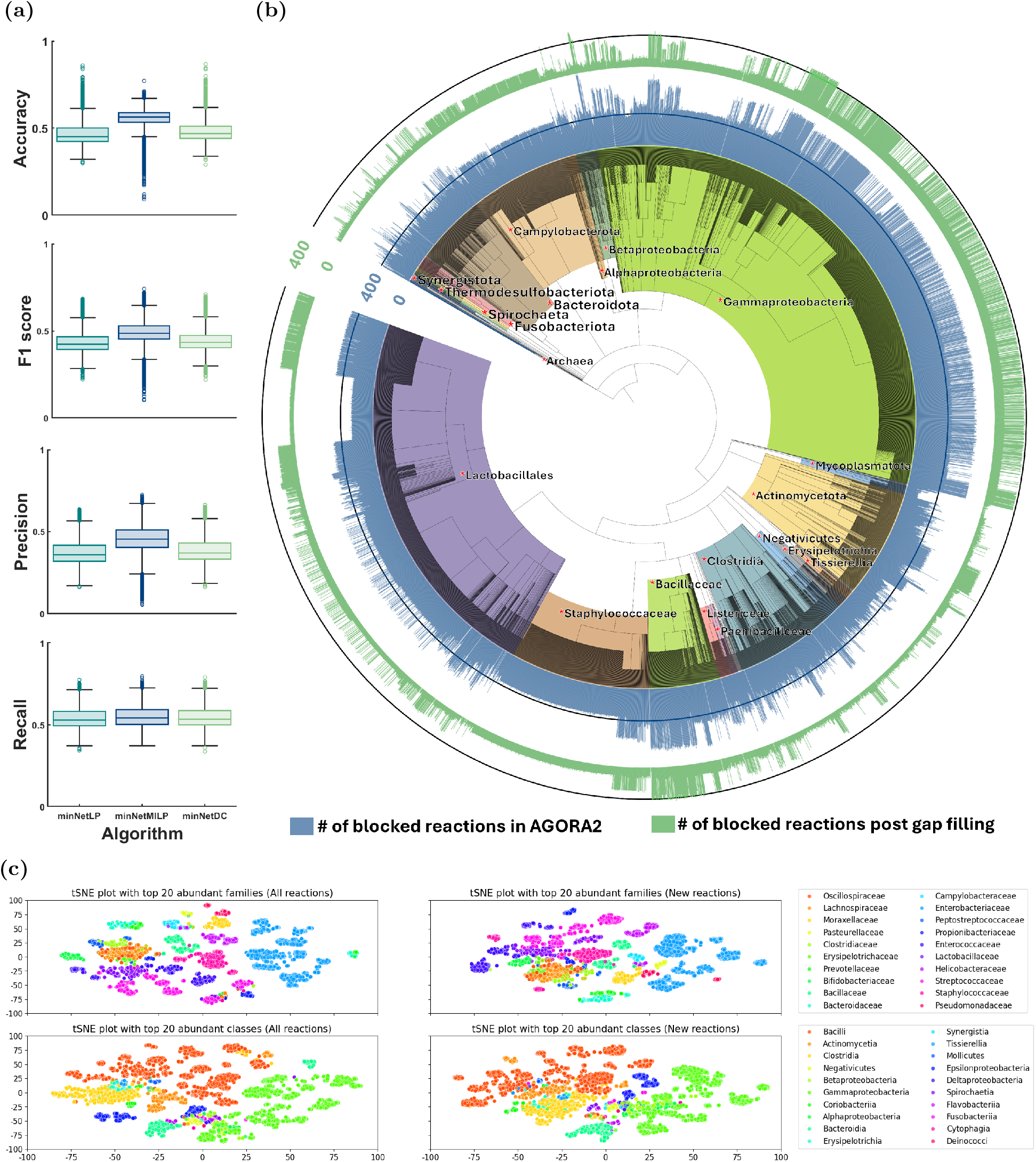
Stoichiometric consistency based gap filling of AGORA2 database. (3a) *minNetMILP* based network inference renders metabolic models with more accuracy and f1-score (*p −value* = 0; right-tailed Wilcoxon signed-rank test). (3b) The number of blocked reactions reduced in all the metabolic models built using SPECTRA. The circular bar plot showing number of blocked reactions before and after gap-filling of the AGORA2 metabolic models. (3c) tSNE clustering of the reactions in newly built models and the newly added reactions, reveal the taxonomic patterns underlying these models.

Subsequently, *minNetMILP* was employed to generate a gap-filled AGORA2 database. All reactions from the original AGORA2 models were treated as core reactions and provided as input to *SPECTRA_ME*. Comparison of the number of blocked reactions before and after gap filling is shown in Fig. 3b. Across all models, at least 58 previously blocked reactions were unblocked, reducing the average number of blocked reactions to approximately 169. The resulting refined models have been publicly deposited and represent a valuable resource for the microbiome research community.

To evaluate the biological relevance of the gap-filled models, t-distributed Stochastic Neighbor Embedding (tSNE) was performed for clustering based on: (i) all reactions in the newly built models (original plus newly added reactions), and (ii) only the newly added reactions. For this, a binary matrix was created indicating the presence (1) or absence (0) of the corresponding reaction across the models. Taxonomic clustering was carried out considering the top 20 abundant classes (7,259 models) and families (6,008 models). The resulting clusters successfully distinguished taxonomic differences between models, demonstrating that SPECTRA preserves phylogenetically meaningful organization. Critically, the clustering patterns observed in newly added reactions which closely mirrored the clustering of all reactions—indicate that SPECTRA systematically identifies and incorporates reactions that are taxonomically relevant across the microbiota. This pattern suggests that gap-filled reactions are not randomly distributed but rather reflect true biological differences in metabolic capabilities across taxa, validating SPECTRA’s ability to recover biologically coherent metabolic improvements. Distinct clusters were obtained by varying the perplexity values of tSNE-based clustering (Supplementary Figs. S12, S13), and all clusters exhibited consistent patterns, further supporting the robustness of these taxonomic distinctions.Additionally, tSNE clustering of subsystems in the gap-filled models revealed similar taxonomic patterns (Supplementary Fig. S14), confirming that SPECTRA’s improvements maintain biological coherence at the functional pathway level. These gap-filled AGORA2 models provide a valuable resource for the microbial modeling community.

Further, the biological relevance of newly added reactions was evaluated using the CLEAN model [45], a machine learning approach trained via contrastive learning to predict EC numbers from protein sequences. Protein sequences from 6,488 of the 7,302 gap-filled models were analyzed. On average, 20.36% of the newly added reactions had predicted EC numbers matching enzymes encoded in the corresponding microbial genomes, demonstrating that these gap-filling predictions are supported by independent protein sequence evidence. This substantial overlap validates SPECTRA’s ability to identify biologically plausible reactions and suggests that machine learning models can effectively prioritize these predictions for experimental validation, bridging constraint-based modeling with modern AI-driven enzyme function prediction. Details on the number of newly added reactions and the exact number of reactions with EC number evidence from CLEAN predictions for each of the models are provided in the Supplementary table S9 Data.

### 2.5 Community-scale gap filling reduces network size and improves metabolic accuracy

GEMs require gap filling step after its draft reconstruction. This step is essential to address incomplete annotations and missing biochemical knowledge, ensuring that the model can sustain growth under defined environmental conditions. Typically, these environmental conditions are implemented as bounds on exchange reactions, which works well for single-organism models. However, in nature, microbes rarely exist in isolation. They coexist within complex communities, exhibiting competition or cooperation depending on the composition of the community and the nutrient environment [32, 46]. Consequently, gap filling for such communities should account for interspecies metabolite exchange and shared resources rather than treating each organism independently.

To evaluate the performance of SPECTRA in this context, we implemented community-scale gap filling on a synthetic human gut microbiota community (hCom) that has been experimentally cultured in defined media [47]. The hCom community comprises 104 microbial members, predominantly from the phyla Firmicutes, Verrucomicrobiota, Bacteroidetes, Actinobacteria, and Proteobacteria. Draft metabolic models for each member were reconstructed using their protein sequences and the CarveMe universal reaction database. SPECTRA_ME was used for the draft network reconstruction with *tradeOff* as the network inference algorithm (refer Methods for more details). We then performed gap filling using both *minNetLP* and *minNetMILP* as network inference algorithms, applying each under two regimes — single-model and community-scale gap filling (refer to Methods 4.10). As shown in Fig. 4a, the number of reactions added by community-scale gap filling is less compared to single-model gap filling for both the network inference algorithms, *minNetLP* and *minNetMILP*, indicating a more parsimonious reconstruction.

**Fig 4.**
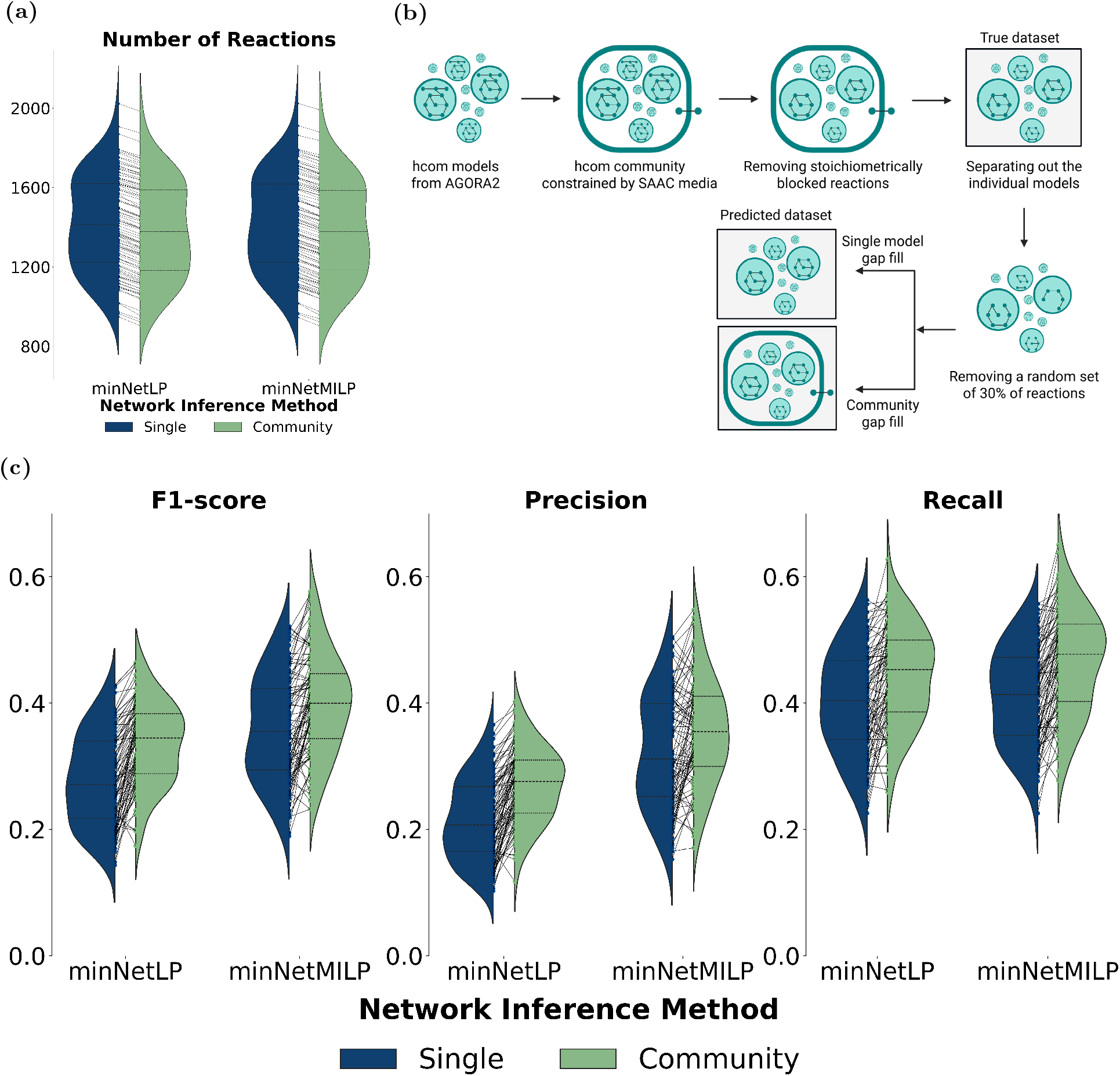
Community scale gap filling generates accurate models with less number of reactions. (4a) The number of reactions added using community-scale gap filling is significantly lower compared to single-model gap filling when using both *minNetLP* and *minNetMILP* as network inference algorithms (*p*-value *≈* 0; left-tailed Wilcoxon signed-rank test). (4b) Overview of the workflow used to compare community-scale and single-model gap filling strategies. (4c) Community-scale gap filling more accurately reconstructs metabolic models than single model gap filling, achieving significantly higher F1-scores, precision, and recall across both *minNetLP* and *minNetMILP* methods (*p*-value *≤* 10^*−*4^; right-tailed Wilcoxon signed-rank test).

Further, to evaluate the accuracy of community-scale gap filling, we used the consistent community model reconstructed from the AGORA2 database under standard amino acid complete (SAAC) medium conditions as the reference dataset (see Methods 4.10). This reference model served as the ground truth against which the gap-filled models obtained using both single-model and community-scale formulations were compared (Fig. 4b). The results presented in Fig. 4c clearly demonstrate the superiority of community-scale gap filling using SPECTRA. Across both network inference algorithms—*minNetLP* and *minNetMILP* —the community-scale approach consistently achieved higher precision, recall, and F1-scores relative to single-model gap filling (*p*-value *≤* 10^*−*4^; right-tailed Wilcoxon signed-rank test). Moreover, the number of reactions added using community-scale gap filling was significantly lower compared to single-model reconstructions (*p*-value *≈* 0; left-tailed Wilcoxon signed-rank test), indicating that the inferred networks are not only more accurate but also more parsimonious. Together, these results highlight SPECTRA’s scalability and its ability to generate biologically meaningful and compact metabolic networks that capture interspecies metabolic dependencies at the community level.

### 2.6 SPECTRA improves recovery of cancer hallmark reactions in cancer metabolic models

We compared the performance of SPECTRA, Fastcore, and Swiftcore in extracting context-specific reactions from a large-scale cancer gene expression dataset. Specifically, transcriptomic data from 1,479 cell lines in the Cancer Cell Line Encyclopedia (CCLE) [48] were used to reconstruct CSMs (see Methods for details). The LocalGini thresholding algorithm [49] was employed to identify core reactions and to assign weights to non-core reactions for each cell line. These inputs formed the basis for CSM reconstruction in all three algorithms. The flux-consistent human GEM, Recon3D, was further improved through the curation of additional reactions and updated gene–protein–reaction (GPR) rules. Details of the development of this improved consensus reconstruction, termed Recon3D+, are provided in Supplementary Texts S1.8 and S1.8.1. The Recon3D+ model is stoichiometrically consistent, contains no blocked reactions, and includes a higher number of reactions, metabolites, and genes compared to the flux-consistent version of Recon3D (Fig. 5a, 5b).

**Fig 5.**
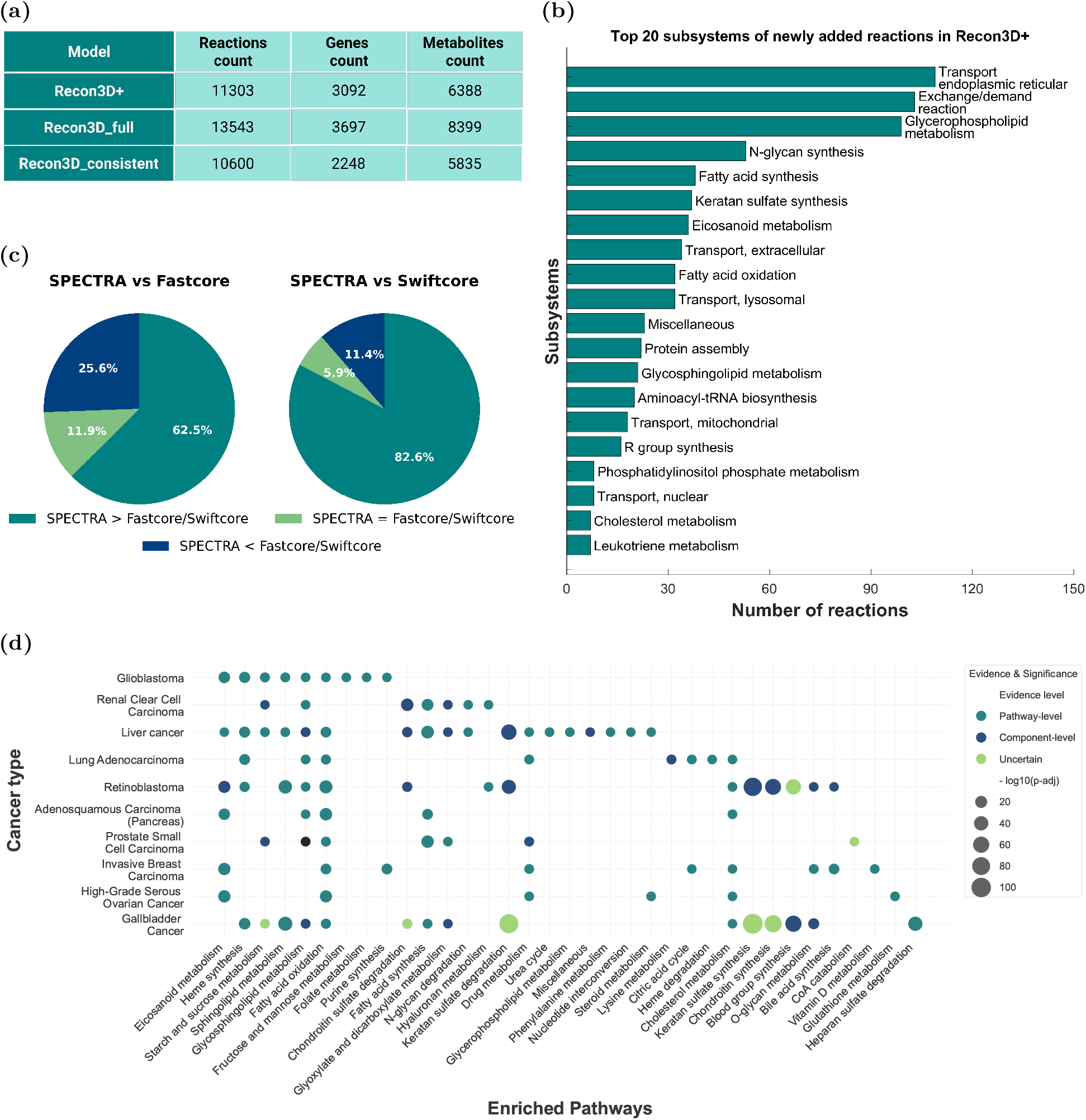
SPECTRA enhances recovery of cancer-relevant metabolic functions across cancer models: (5a) Comparison of reaction, gene, and metabolite counts across different Recon3D models. (5b) Top twenty subsystems ranked by the count of newly added reactions in the Recon3D+ model. (5c) SPECTRA outperformed Fastcore and Swiftcore by capturing a larger number of cancer hallmark reactions, demonstrating its ability to recover biologically relevant, context-specific metabolism in cancer cell lines. The values in the pie chart indicate the percentage of models in which SPECTRA captured a greater than, equal to, or fewer number of hallmark reactions compared with the other methods. (5d) Meta-analysis of enriched metabolic pathways across ten cancer types: The bubble plot illustrates the significance and literature evidence levels for enriched metabolic pathways across various malignancies. The size/color of the bubbles represents the statistical significance indicated by *−log*10(*p −adj*). Pathways are categorized by evidence levels: Pathway-level (high corroboration), Component-level, and Uncertain.

As the universal model, Recon3D+ is same for all the 1,479 models, SPECTRA_ME is used in this case with *minNetLP* as the network inference algorithm. To evaluate performance, we used a curated list of 135 metabolic reactions associated with cancer-specific reprogramming, comprising mainly regulators of the Warburg effect and proliferation, as described in [50]. For each of the 1,479 reconstructed models, we quantified the number of these 135 hallmark reactions recovered and compared the results across the three algorithms (Fig. 5c). As shown, SPECTRA consistently outperformed Fastcore and Swiftcore in capturing a larger number of the cancer-hallmark reactions in 62.5% and 82.6% of the CSMs, respectively, demonstrating its ability to recover biologically relevant context-specific metabolism in cancer cell lines.

As an additional analysis, the predictions from SPECTRA-derived CSMs were validated using published literature. Comparative analysis was performed for the flux prediction of cancer cell line CSMs with their non-cancerous counterparts. The CCLE database includes transcriptomic data for non-cancerous cell lines spanning 10 tissue types: brain, lung, liver, pancreas, kidney, eye, breast, ovary, prostate, and biliary tract. For each of these contexts, Flux Variability Analysis (FVA) [41] was performed on the cancer and the respective non-cancerous models to assess differences in allowable flux ranges. Reactions were classified as downregulated or upregulated in cancer based on two criteria: 1) A reaction was considered downregulated if its maximum flux in the cancer CSM was lower than in the corresponding non-cancerous CSM and its Flux Span Ratio (FSR) was less than 0.5. 2) Conversely, a reaction was considered upregulated if its maximum flux was higher in the cancer CSM and its FSR exceeded 2. The reactions found to be dysregulated between the cancer and non-cancer contexts are provided in Supplementary Data S4 Data (Upregulated reactions) and S5 Data (Downregulated reactions). Flux enrichment analysis (FEA) was done to get the significantly differentially regulated metabolic subsystems in the 10 cancer CSMs, based on the dysregulated reactions (up- and down-regulated) obtained from the FSR results [35]. We validated the predicted enriched pathways for the 10 cancers using the literature, looking for evidence of metabolic dysregulation at the pathway level or at component levels (e.g., genes, enzymes, or metabolites). Pathways lacking supporting studies were classified as uncertain. The details of dysregulated subsystems along with their significance values across the subsystems are provided in Supplementary Data S6 Data. The adjusted p-values were transformed into enrichment scores using a *−* log_10_ transformation for visualization in the bubble plot. A bubble plot elucidating the enrichment of metabolic pathways in 10 cancers with their enrichment scores and strength of evidence in literature are provided in Fig. 5d. The literature evidence of each pathways in 10 cancers has been provided in Supplementary Tables S5 to S14.

Furthermore, from Fig. 5d, it can be seen that there is strong corroboration between the literature findings and in silico predictions for most enriched pathways across many cancers, including glioblastoma, breast cancer, ovarian cancer, pancreatic cancer, lung cancer, liver cancer, and renal clear cell carcinoma. Prostate cancer and retinoblastoma have many component-level pathways compared to complete-pathway level evidence. Gallbladder cancer has many uncertain pathways, which could be attributed to the limited experimental studies on metabolic reprogramming available for it, compared to other cancers discussed [51].

Pathways like fatty acid oxidation and biosynthesis, cholesterol metabolism, glycosphingolipid metabolism, heme synthesis and eicosanoid metabolism, are dysregulated in at least five cancers compared to normal tissues, and they also have adequate pathway-level evidence in many cancers discussed, as seen in Fig. 5d. These enriched metabolic pathways, except heme, represent a lipid-associated metabolic signature in the 10 solid cancers [52]. All these enriched pathways represent a shared pan-cancer signature and have a crucial role in cancer differentiation and progression. Fatty acids and cholesterol support energy production, redox metabolism, and membrane biosynthesis, processes that support cancer development [53–55]. Glycosphingolipids are involved in signalling and epithelial-to-mesenchymal (EMT) transition during metastasis [56]. Heme is a cofactor involved in mitochondrial energy production and modulates signalling pathways that promote tumour growth [57]. Eicosanoids are signalling lipids that regulate tumour growth both positively and negatively by promoting inflammation and immunosuppression [58].

From Fig. 5d, it can be seen that metabolic pathways like chondroitin and keratan metabolisms, and blood group synthesis are not very well explored or studied in many cancers. In cancers, they regulate glycocalyx and extracellular matrix remodelling, which contribute to EMT and altered signalling [59, 60]. More research is required to understand their roles as prognostic markers and for pharmacological targeting in less explored cancers like retinoblastoma and gallbladder cancer. The pathway enrichment analysis with literature validation analysis clearly illustrates the ability of SPECTRA to extract biologically relevant, context-specific reactions.

## 3 Discussion

SPECTRA represents a significant advancement in metabolic model reconstruction by unifying diverse optimization strategies under a single scalable platform. The framework addresses two critical bottlenecks in current constraint-based modeling: the lack of scalable methods for integrating multi-layer omics data and the rigidity of existing reconstruction pipelines. By providing distinct optimization formulations that can accommodate various biological assumptions and constraints, SPECTRA offers unprecedented flexibility compared to specialized tools that are tailored for specific model types or objectives. The scalability of SPECTRA is particularly evident in its application to large-scale model building and community-level modeling. While traditional approaches struggle to integrate collective information from multicellular datasets, SPECTRA’s unified framework can extract metabolic networks ranging from minimal networks to complex multi-cellular models. This scalability is achieved through its modular design, where network connectivity (stoichiometry vs. topology), alternate solution generation (pathway exclusion vs. core reaction directionality), and optimization objectives can be customized based on the biological context.

Our benchmarking results demonstrate that SPECTRA achieves substantial computational improvements over existing state-of-the-art methods. *SPECTRA_ME* consistently outperformed Fastcore and Swiftcore in terms of LP operations while maintaining comparable or superior runtime performance. The 56× speed improvement over FastCC+Fastcore and 36× improvement over SwiftCC+Swiftcore for inconsistent universal reconstructions represents a transformative advance for large-scale metabolic modeling applications. These efficiency gains stem from two key algorithmic innovations. First, *SPECTRA_ME* employs *LPforward* with random-weighted objective function, eliminating the need for repeated LP solves as required by Fastcore (Supplementary text S1.1, S1.2). Second, the inclusion of *LPreverseCC* enables consistency checking in the reverse direction without explicitly changing the signs of columns of the stoichiometric matrix, a computational overhead incurred by Fastcore. These efficiency gains are particularly significant for the metabolic modeling community, where computational bottlenecks often limit the scale and scope of analyses. The ability to simultaneously perform consistency checking and model extraction through *SPECTRA_CC_ME* eliminates the traditional two-step pipeline requirement, reducing the computational overhead. Moreover, since core reactions are always included in the final model, the LP simulations that generates the flux vector, ***v***^*C*^ reduce the number of integer decision variables and constraints in the MILP-based network inference algorithms (*tradeOff* and *minNetMILP* ), further accelerating model reconstruction (Supplementary Fig. S5).

Blocked reaction identification shows the same pattern. *SPECTRA_CC* was the fastest of the four methods tested at every network size above 5,000 reactions, and its LP count did not grow with the size of the network: 6–8 LPs across the full range of single genome-scale models, against up to 3,648 for FastCC and up to 116 for FASTERCC. On expanded Recon3D reconstructions, which represent multicellular systems, the count settled at exactly 10 and remained there from 93,020 to 454,620 reactions, a size at which FASTERCC no longer completes. The practical consequence is that consistency checking need no longer be the bottleneck in reconstruction pipelines for multicellular systems, where it has previously constrained the size of model that could be considered.

Adjudicating by FVA the few reactions on which the methods disagreed also separated two kinds of error. Those made by *SPECTRA_CC* and FASTERCC were numerical and disappeared when the solver tolerance was tightened, whereas those made by SwiftCC persisted, reflecting that it enforces only thermodynamic constraints and so cannot see bounds that block a reaction. A related caveat applies to models whose constraints force reactions to carry flux, such as whole-body reconstructions: relaxing the bounds so that the zero flux vector is feasible, before the consistency check is applied, is what makes the result stable under iteration. When the bounds are not relaxed to include the zero vector solution, removing the identified blocked reactions also removes the reactions that are coupled to these reactions. Hence, the trimmed network admits no feasible flux distribution at all. On the other hand, relaxation of the bounds leads to convergence of a consistent set after a second pass.

SPECTRA’s application to the AGORA2 database demonstrates its utility for large-scale metabolic modeling. The successful gap-filling of over 7,300 metabolic reconstructions with subsequent reduction of blocked reactions from an average of 482 to approximately 169 per model represents a significant resource for the metabolic modeling community. The taxonomic clustering analysis of the gap-filled AGORA2 models revealed meaningful clustering patterns that validate SPECTRA’s ability to preserve phylogenetic relationships while improving model functionality. tSNE visualization of both the complete gap-filled models and specifically the newly added reactions demonstrated clear taxonomic clustering at both family and class levels, indicating that SPECTRA systematically incorporates taxonomically relevant metabolic capabilities. Beyond taxonomic consistency, the integration of the CLEAN model [45] provides independent biological validation for these computational predictions. Analyzing protein sequences from 6,488 models revealed that an average of 20.36% of the newly added reactions align with enzymes encoded directly in the corresponding microbial genomes. This evidence confirms that SPECTRA identifies biologically plausible reactions rather than arbitrary mathematical artifacts. The precise matching of predicted EC numbers to genomic data successfully bridges traditional constraint-based modeling with modern artificial intelligence. Consequently, researchers can use these machine learning predictions to prioritize specific gap-filled reactions for experimental validation. This synergy between mathematical optimization and sequence-level AI effectively narrows the search space for discovering novel enzyme functions.

The most significant contribution of SPECTRA is its profound scalability, which enables modeling of complex, multicellular systems and community-level metabolic networks. While traditional methods reconstruct organisms in isolation, SPECTRA can perform community-scale model reconstruction that accounts for the collective metabolic potential of an entire ecosystem. Our analysis of a synthetic gut microbiota demonstrated that this community-aware approach yields models that are not only more accurate but also more parsimonious, adding significantly fewer reactions than single-model approaches. By capturing interspecies metabolic dependencies, SPECTRA generates reconstructions that better reflect the cooperative and competitive dynamics of real-world microbial communities, a crucial advancement for microbiome research. Not only is SPECTRA extremely scalable, but the reconstructed networks also demonstrate superior coverage of biological relevance (Supplementary text S1.3).

The validation of SPECTRA-derived context-specific models using cancer cell lines provides compelling evidence for the framework’s biological relevance. SPECTRA consistently captured a larger number of cancer-specific hallmark reactions compared to existing methods, demonstrating superior performance in recovering biologically relevant context-specific metabolism. Dysregulated reactions in 10 cancer cell line CSMs obtained by comparing the flux spans with respective non-cancerous cell line CSMs provided enriched metabolic pathways in 10 cancers. A shared pan-cancer metabolic signature was characterised from common enriched pathways in at least 5 cancers, highlighting a lipid-redox axis necessary for cancer proliferation. The majority of the enriched metabolic pathways across all 10 cancers have literature evidence. This biological concordance suggests that SPECTRA’s optimization strategies effectively prioritize metabolically relevant pathways over spurious connections that might arise from purely structural considerations. Our findings indicate that more studies on gallbladder and retinoblastoma are needed to elucidate metabolic rewiring in these cancers. Pathways like chondroitin, keratan, glycan metabolism, etc, are under-researched in cancers and must be investigated for their roles in therapy.

SPECTRA’s versatility extends across multiple applications not fully explored in this study. The framework can be employed for both draft model reconstruction and in gap-filling. For minimal microbiome design, SPECTRA can be configured by assigning non-zero weights exclusively to biomass reactions and incorporating coupling constraints—linear constraints that enforce flux through metabolic reactions only when biomass production is positive. In this configuration, formulations such as *minNetLP* or *minNetMILP* generate parsimonious community models representing the minimal microbial requirements for the required functionalities. SPECTRA’s modular architecture enables applications yet to be fully developed; for example, topological consistency checking identifies reactions blocked solely due to the inability to produce required metabolites, a distinction not captured by existing methods that conflate production and consumption constraints. Future work would address the design of universal models for diverse biological contexts. The community reconstruction approach uses repetitions of bacterial universal models with omics-derived core reactions and weights. This methodology can be extended to host-microbe interactions by integrating species-specific universal models with corresponding omics constraints. Additionally, integration of spatial and single-cell omics data through custom universal models that account for spatial neighborhoods and metabolite diffusion based on distance from blood vessels could substantially improve model accuracy and biological realism. These spatial constraints would enable SPECTRA to generate more accurate reconstructions that reflect the heterogeneous metabolic environments within tissues and organs.

Despite its strengths, SPECTRA has limitations that open avenues for future work. The framework’s performance, particularly for MILP-based formulations like *minNetMILP* and *tradeOff*, can be computationally intensive for extremely large universal reconstructions, such as those for pan-microbiomes. Furthermore, the quality of any extracted model is fundamentally dependent on the accuracy of the underlying universal model and the input omics data. In conclusion, SPECTRA provides a powerful, flexible, and highly efficient platform for metabolic network reconstruction. By unifying disparate methodologies and enabling seamless scaling from single cells to complex communities, it overcomes many of the practical and conceptual barriers that have constrained the field. We anticipate that SPECTRA will become a cornerstone tool for systems biologists, accelerating the translation of complex multi-omics data into predictive and actionable models of metabolism in health and disease.

## 4 Methods

### 4.1 Background

The constraint-based metabolic model captures the metabolic network of *m* metabolites and *n* reactions using the *m × n* stoichiometric matrix ***S***, the reaction set (***R***), the irreversible reaction set (***I ⊆ R***), upper bound (***v***^***ub***^ ***∈*** ℝ^***n***^) and lower bound (***v***^***lb***^ ***∈*** ℝ^***n***^). Each element in the stoichiometric matrix *S*_*ij*_ represents the stoichiometric coefficient of the metabolite *i* in the reaction *j*. The reactions in the metabolic network ***R***, can be categorised as irreversible reactions (***I***) and reversible reactions (***R* \ *I***). Any flux vector (***v*** *∈* ℝ ^*n*^) is said to be a stoichiometrically feasible flux vector if it satisfies the following two conditions.

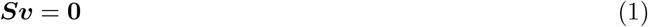

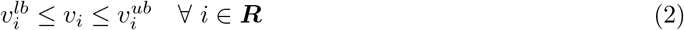

A reaction *i* is said to be stoichiometrically blocked if there exists no stoichiometrically feasible flux vector ***v*** such that *v*_*i*_ ≠ 0. A metabolic model with no stoichiometrically blocked reactions is referred to as a stoichiometrically consistent model.

Similarly, any flux vector (***v****∈* ℝ^*n*^) is said to be a topologically feasible flux vector if it satisfies the following two conditions.

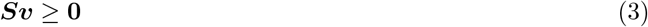

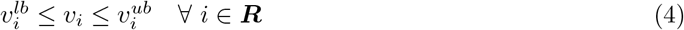

A reaction *i* is said to be topologically blocked if there exists no topologically feasible flux vector ***v*** such that *v*_*i*_ *≠* 0. A metabolic model with no topologically blocked reactions is referred to as a topologically consistent model.

Multi-omics data can be integrated to any universal model using three distinct variables that act as inputs to the SPECTRA class of algorithms: Core reactions (***C*** *∈* ***R***), reaction weights (***w***), and box constraints (***v***^***lb***^ and ***v***^***ub***^). Core reactions are those that are expected to be present in the functional model carrying a non-zero flux in any feasible vector. Reaction weights define the penalties or reaction importance depending the network inference algorithm for the non-core reactions. Box constraints define physiological constraints, such as media composition, minimal biomass production, or ATP maintenance requirements, and must be specified in the universal model. The output of the SPECTRA is a functional model, ***N*** *⊆* ***R***. In next section, we explain distinct optimization formulations used in the study followed the algorithms that use them.

### 4.2 Optimization formulations used in SPECTRA

This section presents eight distinct optimization formulations employed in this study to address the network inference problem. The formulations primarily focus on stoichiometry-based model extraction, with a subsequent discussion on their applicability to topology-based model extraction.

#### 4.2.1 LPforward: To obtain a feasible flux vector from a consistent model with maximum core reactions having a positive flux

| LP problem: LPforward<br>Inputs: $\boldsymbol{S}, \boldsymbol{R}, \boldsymbol{I}, \boldsymbol{v^{ub}}, \boldsymbol{v^{lb}}, \boldsymbol{K}$<br>Outputs: $\boldsymbol{v}$ |  |  |  |
|---|---|---|---|
| $\max_{\boldsymbol{v},\boldsymbol{z}}$ | $\sum_{i \in \boldsymbol{K} \text{ and } i \notin \boldsymbol{I}} x_i.z_i$ |  |  |
| s.t. | $\boldsymbol{Sv} = \boldsymbol{0}$ |  |  |
|  | $v_i^{lb} \leq v_i \leq v_i^{ub}$ | $\forall\, i \in \boldsymbol{R}$ | (LPforward) |
|  | $v_i \geq z_i$ | $\forall\, i \in \boldsymbol{K}$and $i \notin \boldsymbol{I}$ |  |
|  | $z_i \leq \epsilon$ | $\forall\, i \in \boldsymbol{K}$and $i \notin \boldsymbol{I}$ |  |
|  | $v_i \geq \epsilon$ | $\forall\, i \in \boldsymbol{K}$and $i \in \boldsymbol{I}$ |  |

Given a set of reactions ***K*** *⊆* ***R***, the goal of *LPforward* is to maximize the number of reactions in ***K*** that carry a positive flux in a consistent model.

The first two constraints ensure the feasibility of the flux vector. A set of auxiliary decision variables ***z*** is introduced, where each *z*_*i*_ serves as a lower bound for the flux of reversible reactions in ***K***. These variables are bounded above by a small positive constant *ϵ*, enforcing that maximization of *z*_*i*_ promotes activation of reversible reactions with positive flux.

In Equation-(LPforward), the *z*_*i*_ terms are weighted in the objective function by random coefficients *x*_*i*_, to avoid degeneracy and ensure a non-zero objective (see Supplementary Information S1.1 for details). In this study, *x*_*i*_ are sampled from a uniform distribution: *x*_*i*_ *∼* ***U*** (1, 1.1).

Finally, all irreversible reactions in ***K*** are explicitly constrained to carry positive flux. This guarantees that a single run of the formulation in Equation-(LPforward) suffices to activate all irreversible reactions in ***K***.

#### 4.2.2 LPforwardCC: To obtain a feasible flux vector from a model with maximum core reactions having a positive flux

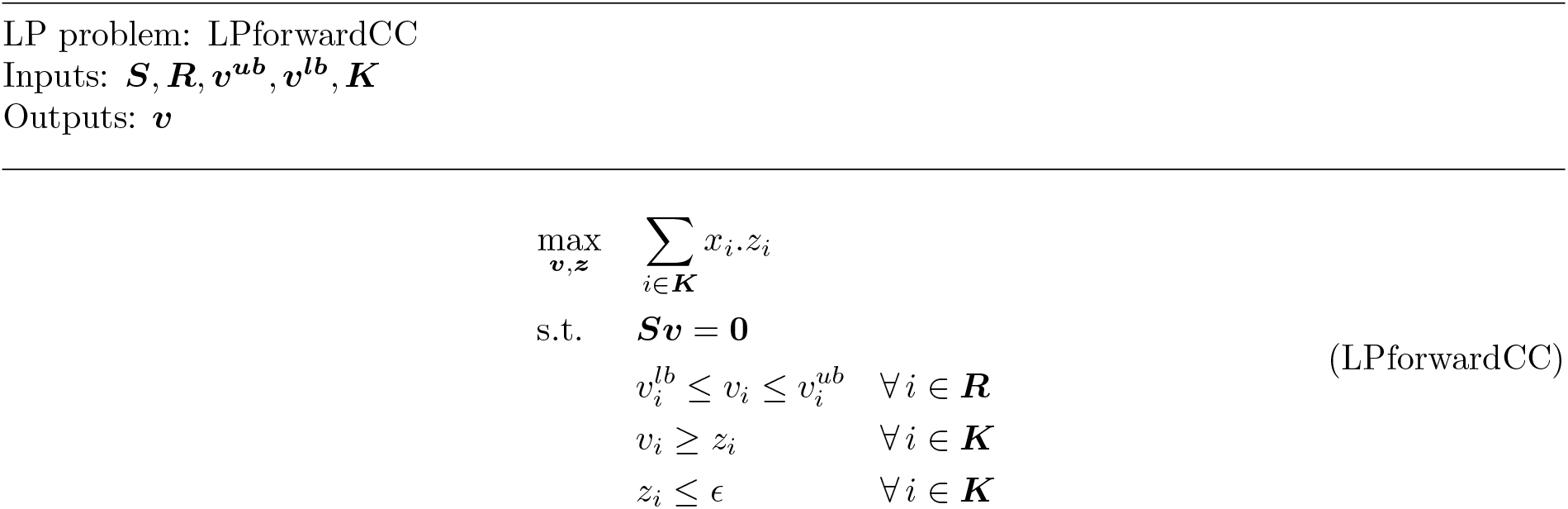

| LP problem: LPforwardCC<br>Inputs: $\boldsymbol{S}, \boldsymbol{R}, \boldsymbol{v}^{\boldsymbol{ub}}, \boldsymbol{v}^{\boldsymbol{lb}}, \boldsymbol{K}$<br>Outputs: $\boldsymbol{v}$ |  |  |  |
|---|---|---|---|
| $\max\limits_{\boldsymbol{v},\boldsymbol{z}}$ | $\sum\limits_{i\in\boldsymbol{K}} x_i.z_i$ |  | (LPforwardCC) |
| s.t. | $\boldsymbol{S}\boldsymbol{v} = \boldsymbol{0}$ |  |  |
|  | $v_i^{lb} \leq v_i \leq v_i^{ub}$ | $\forall\, i \in \boldsymbol{R}$ |  |
|  | $v_i \geq z_i$ | $\forall\, i \in \boldsymbol{K}$ |  |
|  | $z_i \leq \epsilon$ | $\forall\, i \in \boldsymbol{K}$ |  |

The *LPforwardCC* formulation follows the same principles as *LPforward*, but without enforcing positive flux for irreversible reactions in ***K*** (***K*** *∩* ***I***). This modification is necessary because blocked irreversible reactions may render the LP problem infeasible. Thus, given ***K*** *⊆* ***R***, *LPforwardCC* maximizes the number of reactions in ***K*** that carry positive flux in the forward direction, as shown in Equation-(LPforwardCC).

### 4.2.3 LPreverseCC: To obtain a feasible flux vector from a model with maximum core reactions having a negative flux

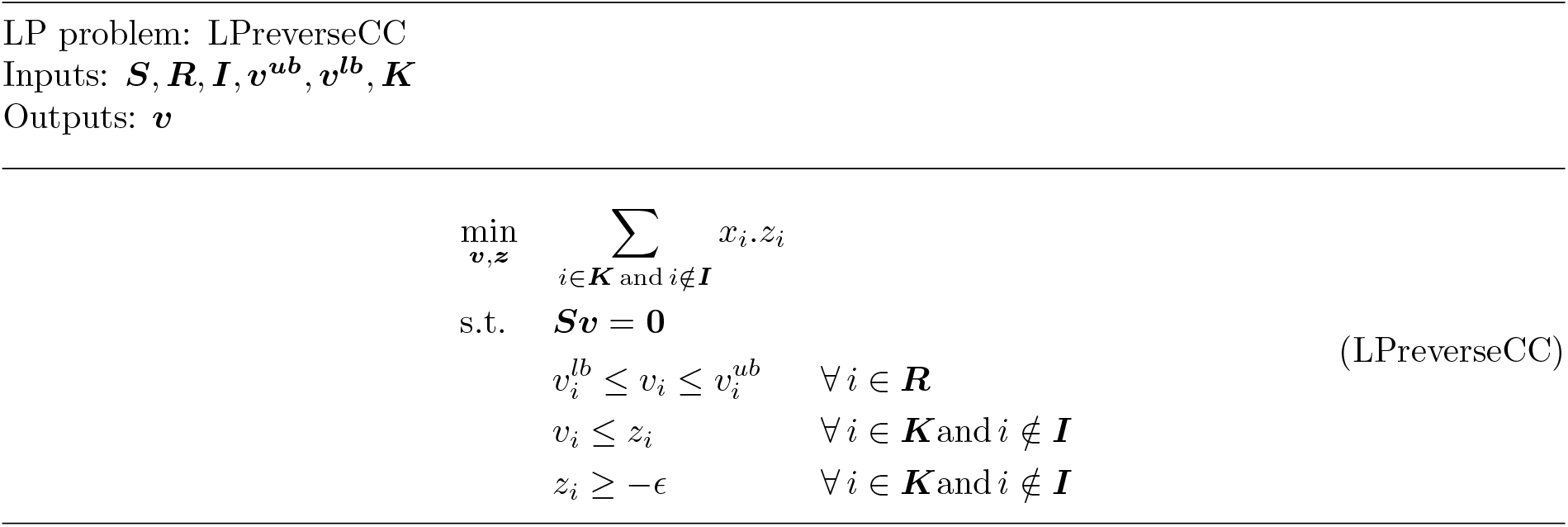

| LP problem: LPreverseCC<br>Inputs: ***S***, ***R***, ***I***, ***v***<sup>***ub***</sup>, ***v***<sup>***lb***</sup>, ***K***<br>Outputs: ***v*** |  |  |  |
|---|---|---|---|
| min<sub>***v***,***z***</sub> | $\sum_{i\in\boldsymbol{K}\text{ and }i\notin\boldsymbol{I}} x_i.z_i$ |  | (LPreverseCC) |
| s.t. | ***Sv*** = **0** |  |  |
|  | *v*<sub>*i*</sub><sup>*lb*</sup> ≤ *v*<sub>*i*</sub> ≤ *v*<sub>*i*</sub><sup>*ub*</sup> | ∀ *i* ∈ ***R*** |  |
|  | *v*<sub>*i*</sub> ≤ *z*<sub>*i*</sub> | ∀ *i* ∈ ***K*** and *i* ∉ ***I*** |  |
|  | *z*<sub>*i*</sub> ≥ −ϵ | ∀ *i* ∈ ***K*** and *i* ∉ ***I*** |  |

The *LPreverseCC* formulation is the exact counterpart of *LPforwardCC*. Given a set of reactions ***K*** *⊆* ***R***, the objective is to maximize the number of reactions in ***K*** that carry flux in the reverse (negative) direction.

The first two constraints ensure the feasibility of the flux vector. A set of auxiliary decision variables ***z*** is introduced, where each *z*_*i*_ serves as an upper bound for the flux of reversible reactions in ***K***. These variables are bounded below by *−ϵ*, such that minimizing *z*_*i*_ promotes activation of reversible reactions with negative flux. The resulting formulation is provided in Equation-(LPreverseCC).

#### 4.2.4 minNetLP: Finding the minimal subnetwork by reducing the absolute sum of fluxes

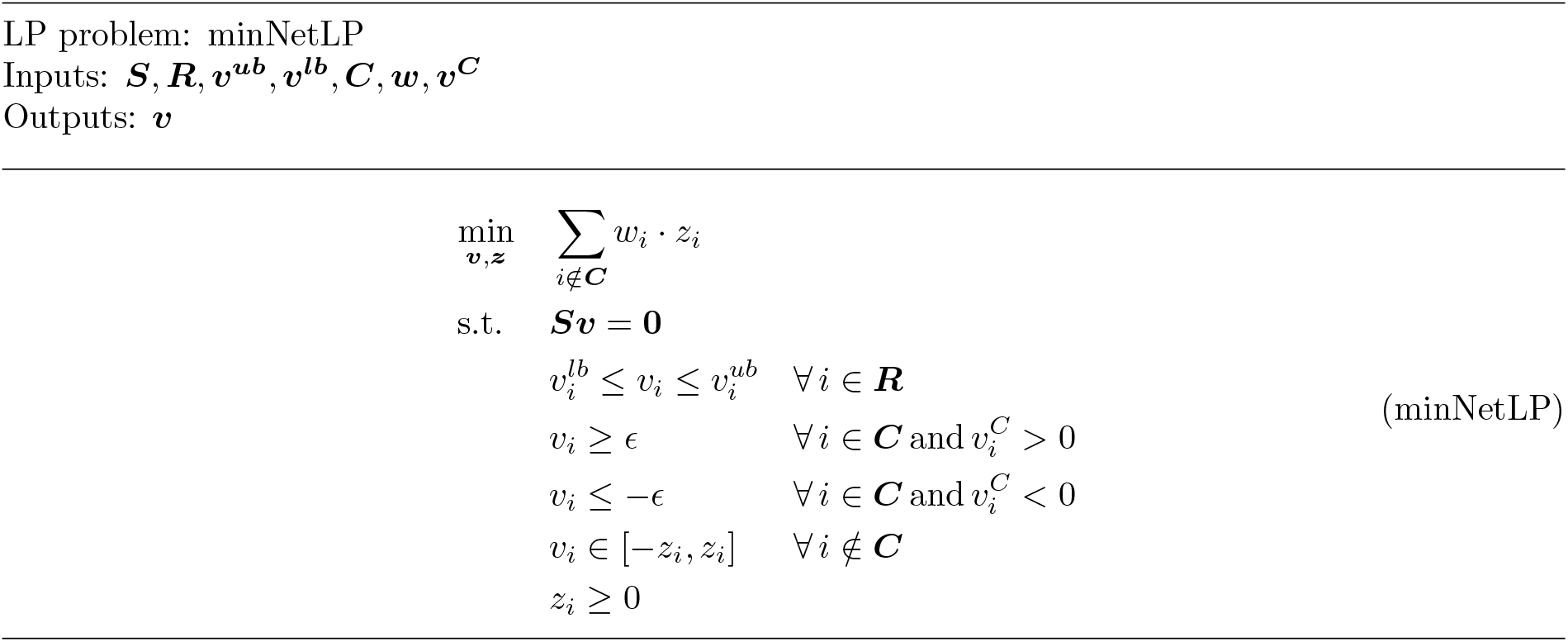

| LP problem: minNetLP<br>Inputs: ***S***, ***R***, ***v***<sup>***ub***</sup>, ***v***<sup>***lb***</sup>, ***C***, ***w***, ***v***<sup>***C***</sup><br>Outputs: ***v*** |  |  |  |
|---|---|---|---|
| $\min_{\boldsymbol{v},\boldsymbol{z}}$ | $\sum_{i \notin \boldsymbol{C}} w_i \cdot z_i$ |  | (minNetLP) |
| s.t. | ***Sv*** = **0** |  |  |
|  | *v*<sub>*i*</sub><sup>*lb*</sup> ≤ *v*<sub>*i*</sub> ≤ *v*<sub>*i*</sub><sup>*ub*</sup> | ∀ *i* ∈ ***R*** |  |
|  | *v*<sub>*i*</sub> ≥ *ϵ* | ∀ *i* ∈ ***C*** and *v*<sub>*i*</sub><sup>*C*</sup> > 0 |  |
|  | *v*<sub>*i*</sub> ≤ −*ϵ* | ∀ *i* ∈ ***C*** and *v*<sub>*i*</sub><sup>*C*</sup> < 0 |  |
|  | *v*<sub>*i*</sub> ∈ [−*z*<sub>*i*</sub>, *z*<sub>*i*</sub>] | ∀ *i* ∉ ***C*** |  |
|  | *z*<sub>*i*</sub> ≥ 0 |  |  |

The objective of the *minNetLP* is to minimize the inclusion of non-core reactions in the final functional model. The linear programming (LP) formulation, *minNetLP*, achieves this by minimizing the total flux through non-core reactions, while ensuring that all core reactions carry non-zero flux.

Given a universal reconstruction (***S, R, v***^***ub***^, ***v***^***lb***^), a set of core reactions ***C*** *⊆* ***R***, reaction weights ***w***, and a feasible flux vector ***v***^***C***^ such that 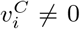 for all *i ∈* ***C***, *minNetLP* solves a linear program to identify a flux vector ***v*** that is dense in core reactions and sparse in non-core reactions. The formulation is given in Equation-(minNetLP). The method to obtain feasible flux vector, ***v***^***C***^ will be explained in section 4.3.

The number of decision variables in *LPminimal* is |***R***| + |***R*** *\* ***C***|. This formulation introduces an auxiliary variable *z*_*i*_ to minimize the *ℓ*_1_-norm of the flux through each non-core reaction *v*_*i*_ [61]. The weights *w*_*i*_ allow for adjusting the relative penalty associated with including each non-core reaction. Unless otherwise specified, all non-core reactions are assigned equal weights (*w*_*i*_ = 1). The first two constraints ensure that the resulting flux vector is stoichiometrically feasible and lies within the specified flux bounds. The next set of constraints enforces a non-zero flux through all core reactions in ***C***. The parameter *ϵ* denotes the minimum flux required for a reaction to be considered active. After solving the LP, reactions with absolute flux values greater than zero are included in the final functional model, ***N*** *⊆* ***R***.

#### 4.2.5 minNetMILP: Finding the minimal subnetwork by reducing the number of reactions

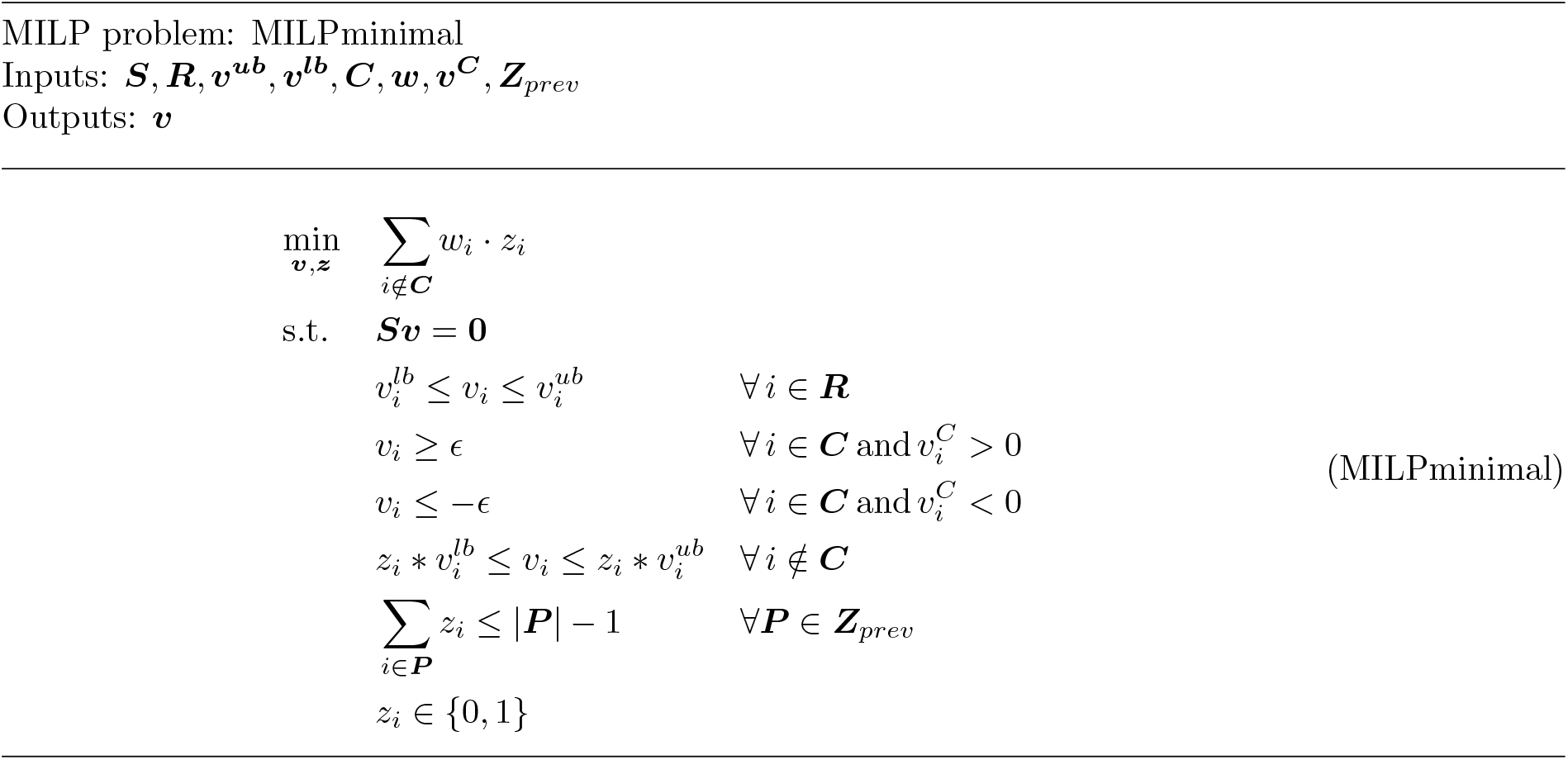

| MILP problem: MILPminimal<br>Inputs: $\boldsymbol{S}, \boldsymbol{R}, \boldsymbol{v^{ub}}, \boldsymbol{v^{lb}}, \boldsymbol{C}, \boldsymbol{w}, \boldsymbol{v^{C}}, \boldsymbol{Z}_{prev}$<br>Outputs: $\boldsymbol{v}$ |  |  |  |
|---|---|---|---|
| $\min_{\boldsymbol{v},\boldsymbol{z}}$ | $\sum_{i \notin \boldsymbol{C}} w_i \cdot z_i$ |  |  |
| s.t. | $\boldsymbol{S}\boldsymbol{v} = \boldsymbol{0}$ |  |  |
|  | $v_i^{lb} \leq v_i \leq v_i^{ub}$ | $\forall\, i \in \boldsymbol{R}$ |  |
|  | $v_i \geq \epsilon$ | $\forall\, i \in \boldsymbol{C} \,\text{and}\, v_i^{C} > 0$ | (MILPminimal) |
|  | $v_i \leq -\epsilon$ | $\forall\, i \in \boldsymbol{C} \,\text{and}\, v_i^{C} < 0$ |  |
|  | $z_i * v_i^{lb} \leq v_i \leq z_i * v_i^{ub}$ | $\forall\, i \notin \boldsymbol{C}$ |  |
|  | $\sum_{i \in \boldsymbol{P}} z_i \leq \|\boldsymbol{P}\| - 1$ | $\forall \boldsymbol{P} \in \boldsymbol{Z}_{prev}$ |  |
|  | $z_i \in \{0, 1\}$ |  |  |

An alternative formulation, *minNetMILP*, minimizes the number of non-core reactions included in the final model by posing the problem as a mixed-integer linear program (MILP). The formulation is given in Equation-(MILPminimal). All variables retain the same definitions as in *minNetLP* .

In this formulation, *z*_*i*_ is a binary decision variable that indicates whether the non-core reaction *i* is included (*z*_*i*_ = 1) or excluded (*z*_*i*_ = 0) from the final model. Also, this formulation takes the set of ids of previously obtained solutions as ***Z***_*prev*_ in order to get an alternate solution that is not previously obtained. After solving the MILP, reactions with absolute flux values greater than zero are included in the final functional model, ***N*** *⊆* ***R***.

#### 4.2.6 tradeOff: Finding the pathways with a net positive weights

The objective of *tradeOff* is to construct a consistent subnetwork that balances the inclusion of positively weighted reactions and the exclusion of negatively weighted reactions, as formulated in Equation-(tradeOff).

Given a universal reconstruction (***S, R, I, v***^***ub***^, ***v***^***lb***^), a set of core reactions ***C*** *⊆* ***R***, reaction weights ***w***, and a feasible flux vector ***v***^***C***^ such that 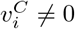 for all *i ∈****C***, *tradeOff* solves for a flux vector ***v*** together with a set of binary variables governing the inclusion of non-core reactions. For each non-core reaction, a binary variable *z*_*i*_ is introduced, taking value 1 if the reaction is included in the network and 0 otherwise. The objective maximizes the weighted sum of these binary variables, thereby favoring reactions with positive weights while penalizing negatively weighted ones. In addition, two auxiliary binary variables, *a*_*i*_ and *b*_*i*_, are introduced for every non-core reversible reaction to ensure directionality consistency: a reversible reaction may be active in either the forward or reverse direction, but not both simultaneously. After solving the MILP, all reactions with absolute flux values greater than zero are retained to define the final functional subnetwork ***N*** *⊆* ***R***.

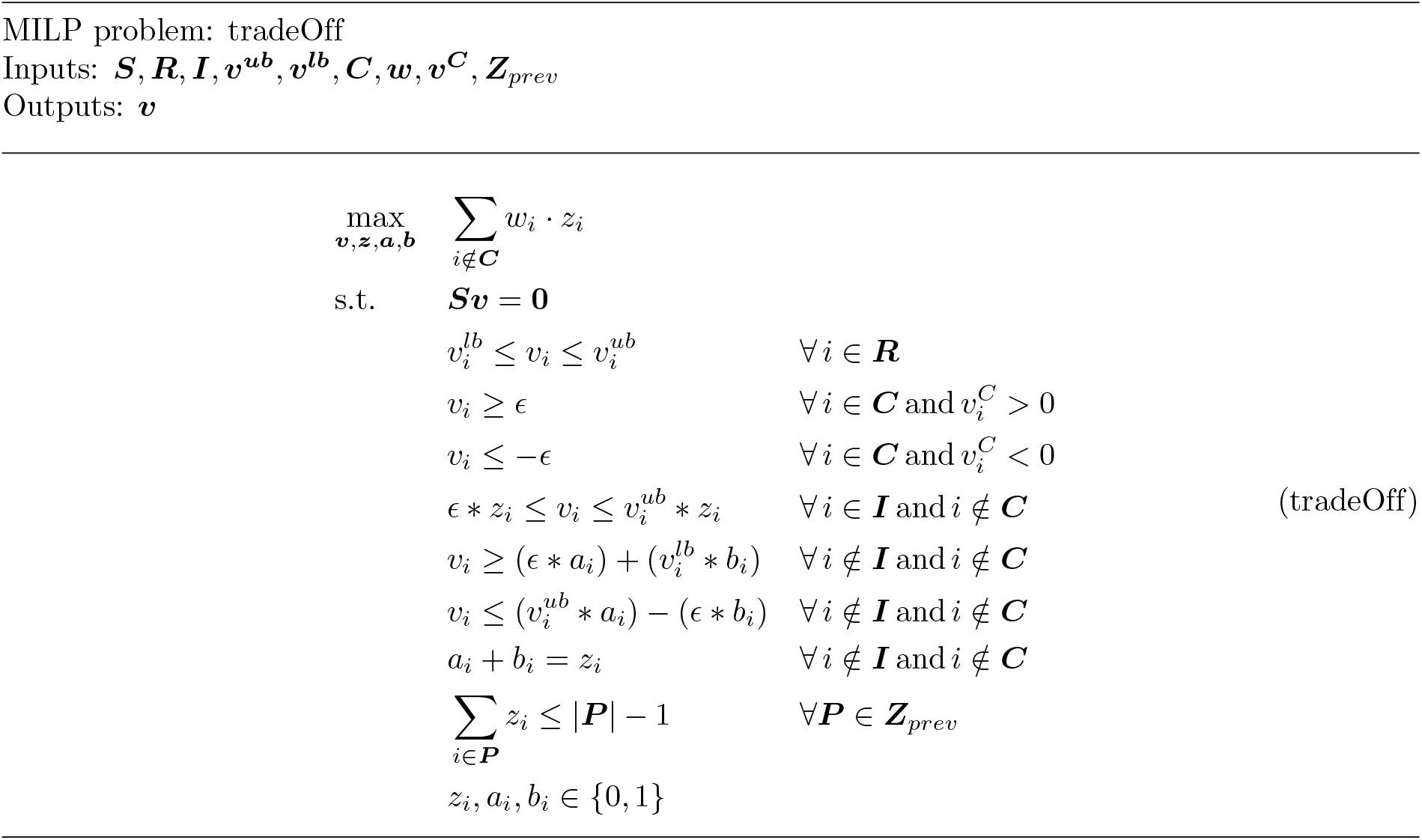

| MILP problem: tradeOff<br>Inputs: **S**, **R**, **I**, **v<sup>ub</sup>**, **v<sup>lb</sup>**, **C**, **w**, **v<sup>C</sup>**, **Z**<sub>prev</sub><br>Outputs: **v** |  |  |  |
|---|---|---|---|
| max<sub>**v**,**z**,**a**,**b**</sub> | ∑<sub>i∉**C**</sub> w<sub>i</sub> · z<sub>i</sub> |  |  |
| s.t. | **Sv** = **0** |  |  |
|  | v<sub>i</sub><sup>lb</sup> ≤ v<sub>i</sub> ≤ v<sub>i</sub><sup>ub</sup> | ∀ i ∈ **R** |  |
|  | v<sub>i</sub> ≥ ϵ | ∀ i ∈ **C** and v<sub>i</sub><sup>C</sup> > 0 |  |
|  | v<sub>i</sub> ≤ −ϵ | ∀ i ∈ **C** and v<sub>i</sub><sup>C</sup> < 0 |  |
|  | ϵ ∗ z<sub>i</sub> ≤ v<sub>i</sub> ≤ v<sub>i</sub><sup>ub</sup> ∗ z<sub>i</sub> | ∀ i ∈ **I** and i ∉ **C** | (tradeOff) |
|  | v<sub>i</sub> ≥ (ϵ ∗ a<sub>i</sub>) + (v<sub>i</sub><sup>lb</sup> ∗ b<sub>i</sub>) | ∀ i ∉ **I** and i ∉ **C** |  |
|  | v<sub>i</sub> ≤ (v<sub>i</sub><sup>ub</sup> ∗ a<sub>i</sub>) − (ϵ ∗ b<sub>i</sub>) | ∀ i ∉ **I** and i ∉ **C** |  |
|  | a<sub>i</sub> + b<sub>i</sub> = z<sub>i</sub> | ∀ i ∉ **I** and i ∉ **C** |  |
|  | ∑<sub>i∈**P**</sub> z<sub>i</sub> ≤ \|**P**\| − 1 | ∀**P** ∈ **Z**<sub>prev</sub> |  |
|  | z<sub>i</sub>, a<sub>i</sub>, b<sub>i</sub> ∈ {0, 1} |  |  |

### 4.2.7 optimBiomass: Finding the subnetwork that optimizes flux through the biomass reaction

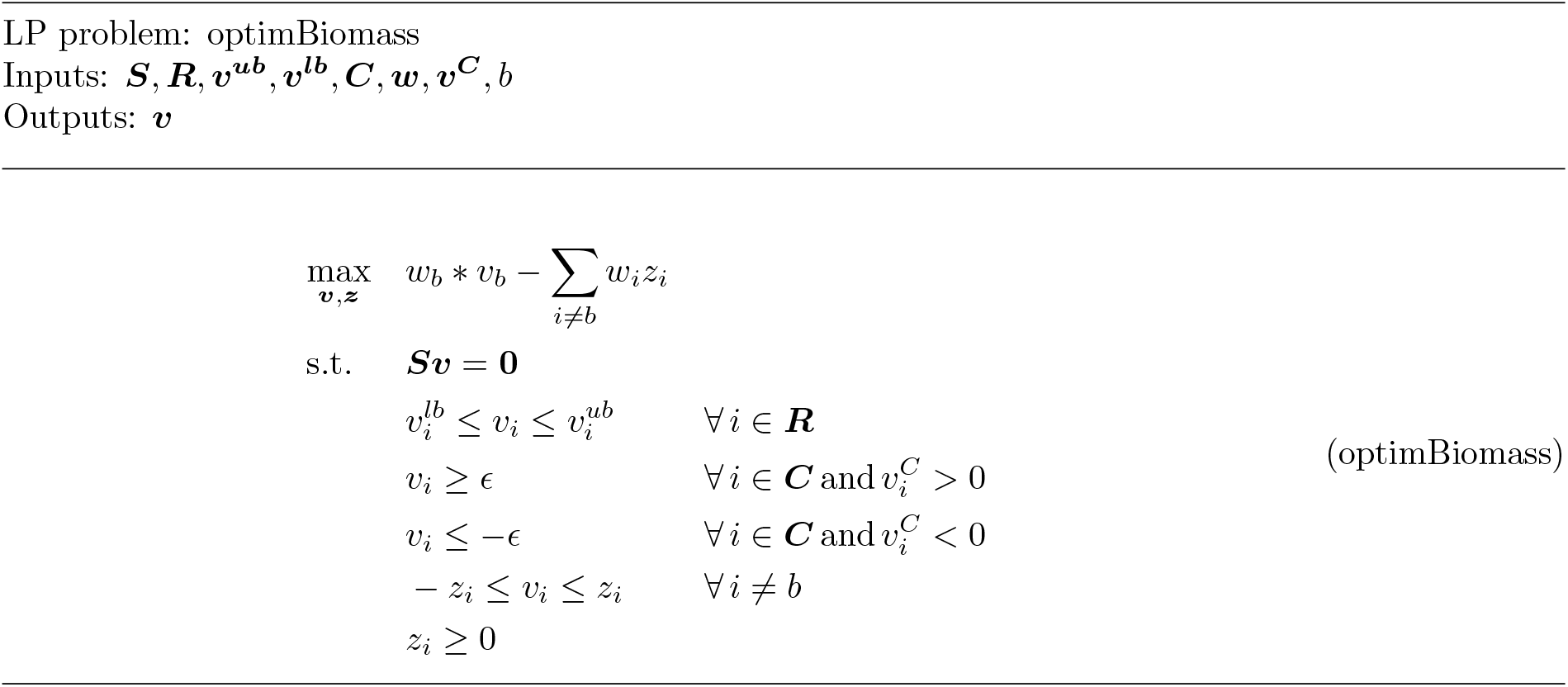

| LP problem: optimBiomass<br>Inputs: $\boldsymbol{S}, \boldsymbol{R}, \boldsymbol{v^{ub}}, \boldsymbol{v^{lb}}, \boldsymbol{C}, \boldsymbol{w}, \boldsymbol{v^{C}}, b$<br>Outputs: $\boldsymbol{v}$ |  |  |  |
|---|---|---|---|
| $\max_{\boldsymbol{v},\boldsymbol{z}}$ | $w_b * v_b - \sum_{i \neq b} w_i z_i$ |  | (optimBiomass) |
| s.t. | $\boldsymbol{S}\boldsymbol{v} = \boldsymbol{0}$ |  |  |
|  | $v_i^{lb} \leq v_i \leq v_i^{ub}$ | $\forall\, i \in \boldsymbol{R}$ |  |
|  | $v_i \geq \epsilon$ | $\forall\, i \in \boldsymbol{C}$ and $v_i^C > 0$ |  |
|  | $v_i \leq -\epsilon$ | $\forall\, i \in \boldsymbol{C}$ and $v_i^C < 0$ |  |
|  | $-z_i \leq v_i \leq z_i$ | $\forall\, i \neq b$ |  |
|  | $z_i \geq 0$ |  |  |

The objective of the LP formulation, *optimBiomass* is to increase the flux through the biomass reactions and minimize the sum of absolute value of the fluxes through all other reactions. The entire formulation is shown in the Equation-(optimBiomass). The variable, *b* denote the biomass reaction index in the input model. All the other variables here have the same meaning as defined in the previous subsections. Note that *b* could be a single index or a vector of indices amenable for multicellular modeling. The weights, *w*_*i*_ defined here work as penalties to all the other reactions and act as reward for the biomass reaction. After solving the LP, all reactions with absolute flux values greater than zero are retained to define the final functional subnetwork ***N*** *⊆* ***R***.

### 4.3 Obtaining a feasible flux vector that has non-zero flux through the core reactions

The network inference optimization formulations require the flux vector, ***V*** ^***C***^ as an input. This vector has non-zero flux through all reactions in ***C***, the flux vectors obtained from *LPforward* (or *LPforwardCC* ) and *LPreverseCC* are iteratively combined using a linear algebraic approach, which is described in this and next section. The following theorem has been used to combine the results from *LPforward* and *LPreverseCC* to get ***v***^***C***^.

#### Theorem 1.

*If* ***v***^**1**^ *and* ***v***^**2**^ *are stoichiometrically feasible flux vectors, then their convex combination is also a stoichiometrically feasible flux vector*.

*Proof*. Any flux vector must satisfy Equations-(1) and (2) to be stoichiometrically feasible.

**For Equation-(1):**

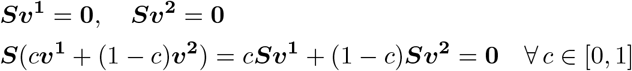

**For Equation-(2):**

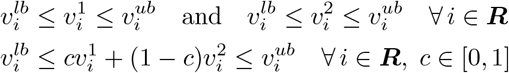

If the universal reconstruction includes only thermodynamic (or irreversibility) constraints, a conic combination of two stoichiometrically feasible flux vectors results in another stoichiometrically feasible flux vector. However, in general, metabolic models incorporate additional constraints such as physicochemical, media, enzyme capacity, and regulatory constraints—all of which are represented through lower and upper bounds on reaction fluxes. In such cases, only a convex combination of stoichiometrically feasible vectors is guaranteed to satisfy these bounds. Accordingly, the optimized flux vectors obtained from *LPforward* and *LPreverseCC* are iteratively combined in a convex manner to construct the final flux vector ***v***^***C***^.

### 4.4 To impose topological constraints for network inference

All optimization formulations described above, as well as Theorem 1, can be adapted to incorporate topological constraints. Specifically for the optimization formulations, this is achieved by replacing the stoichiometric steady-state constraints (***Sv*** = **0**) with topological constraints (***Sv*** *≥* **0**). Furthermore, Theorem 1 remains valid under topological feasibility: the convex combination of two topologically feasible flux vectors that satisfy Equations-(3) and (4) is also topologically feasible.

### 4.5 SPECTRA_ME: Algorithm to extract metabolic networks from a consistent universal reconstruction

*SPECTRA_ME* constructs a consistent metabolic subnetwork in which no reactions are blocked. Blocked reactions may arise from topological or stoichiometric constraints, depending on the structural assumptions (see Section S1.7). The type of structural constraint to impose is defined by the user. Given the mass-balance and box constraints (Equations-(1) and (2)), reactions in a metabolic model are inherently coupled. *SPECTRA_ME* leverages a feasible flux vector, ***v***^***C***^, to encode these coupling relationships among core reactions. The construction of ***v***^***C***^ relies on the formulations in Equations-(LPforward) and (LPreverseCC). Specifically, in each iteration of the while loop (step 3 in Algorithm 1), Equation-(LPforward) identifies the maximal subset of core reactions capable of carrying flux in the forward direction, while Equation-(LPreverseCC) does the same for reversible core reactions in the reverse direction. The resulting flux vectors, ***v***_***f***_ and ***v***_***r***_, are then convexly combined with ***v***^***C***^, as described in Theorem 1.

Irreversible core reactions are excluded from *LPreverseCC*, since they cannot carry negative flux. Once the coupling relationships are captured in ***v***^***C***^, one of the five network inference algorithms is applied via the *netInf* function in Algorithm 1. The choice of algorithm is determined by the parameter *probType*, which can be set to *minNetLP, minNetMILP, minNetDC, tradeOff*, or *optimBiomass*. Because convex combinations of feasible vectors may yield flux values smaller than *ϵ* for some core reactions, the threshold *ϵ* is scaled by a factor of 0.1 when updating the active set ***K***. Finally, note that the universal reconstruction provided as input must itself be consistent, with no blocked reactions present.

#### Algorithm 1

*SPECTRA_ME*: Algorithm to extract metabolic networks from a consistent universal reconstruction

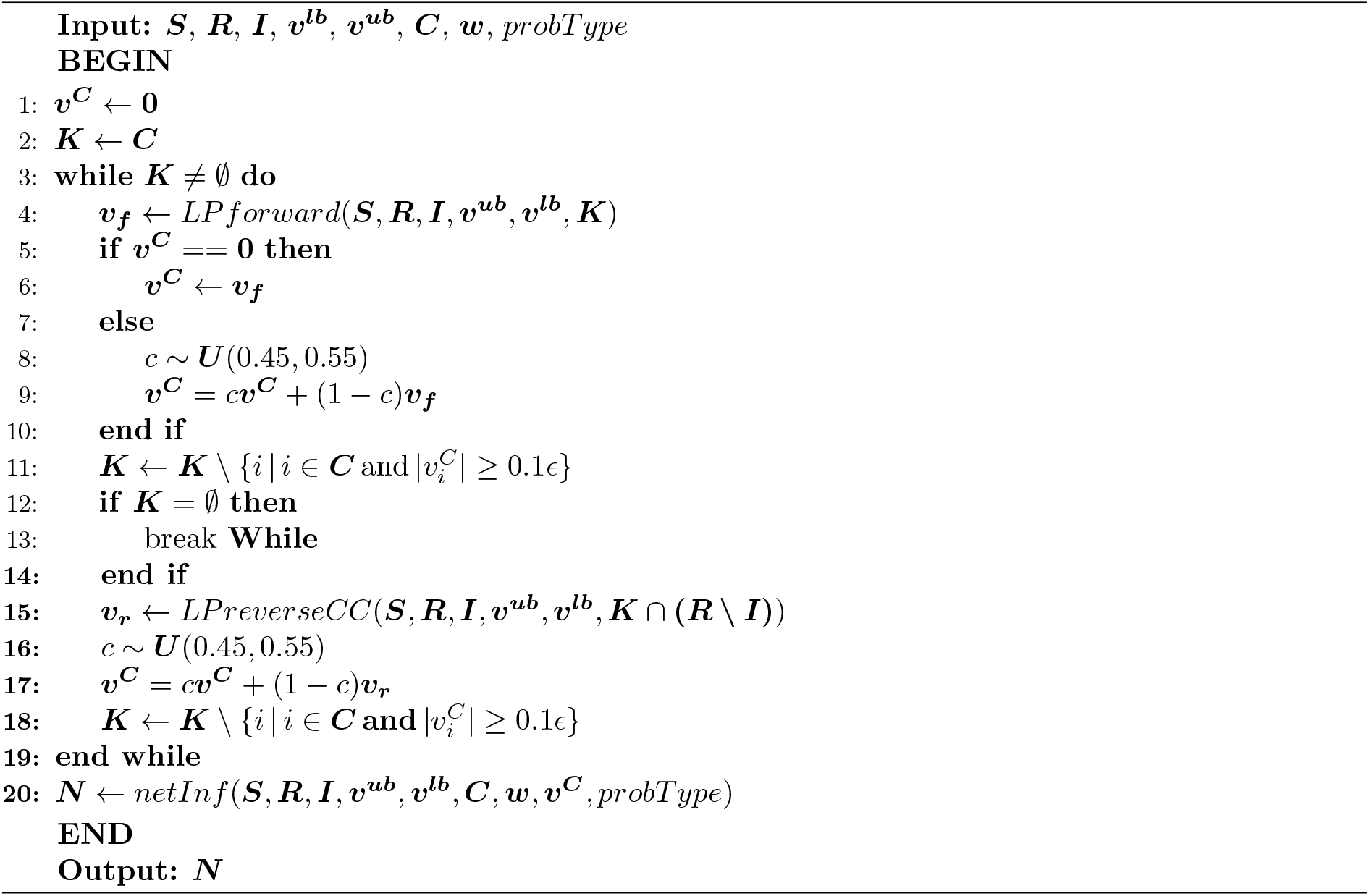

### 4.6 SPECTRA_CC_ME: Algorithm to extract metabolic networks from a universal reconstruction

*SPECTRA_CC_ME* can be viewed as a combination of *SPECTRA_CC* (refer to Methods 4.7) and *SPECTRA_ME*, simultaneously performing the detection and removal of blocked reactions from the universal reconstruction, as well as model extraction. It returns both the extracted model and a list of blocked core reactions. The pseudocode for *SPECTRA_CC_ME* is presented in Algorithm 2. The flux vector ***v***^***C***^ is generated in a manner similar to *SPECTRA_ME*, except that all reactions in the model are treated as core reactions, and *LPforwardCC* is used in place of *LPforward*. The set of blocked core reactions is denoted as ***BlockedCore***. These reactions are removed from the core set, as their presence would render the network inference algorithms infeasible. Subsequently, either of the five network inference algorithms are applied to extract the metabolic network.

#### Algorithm 2

*SPECTRA_CC_ME*: Algorithm to extract metabolic networks from a universal reconstruction

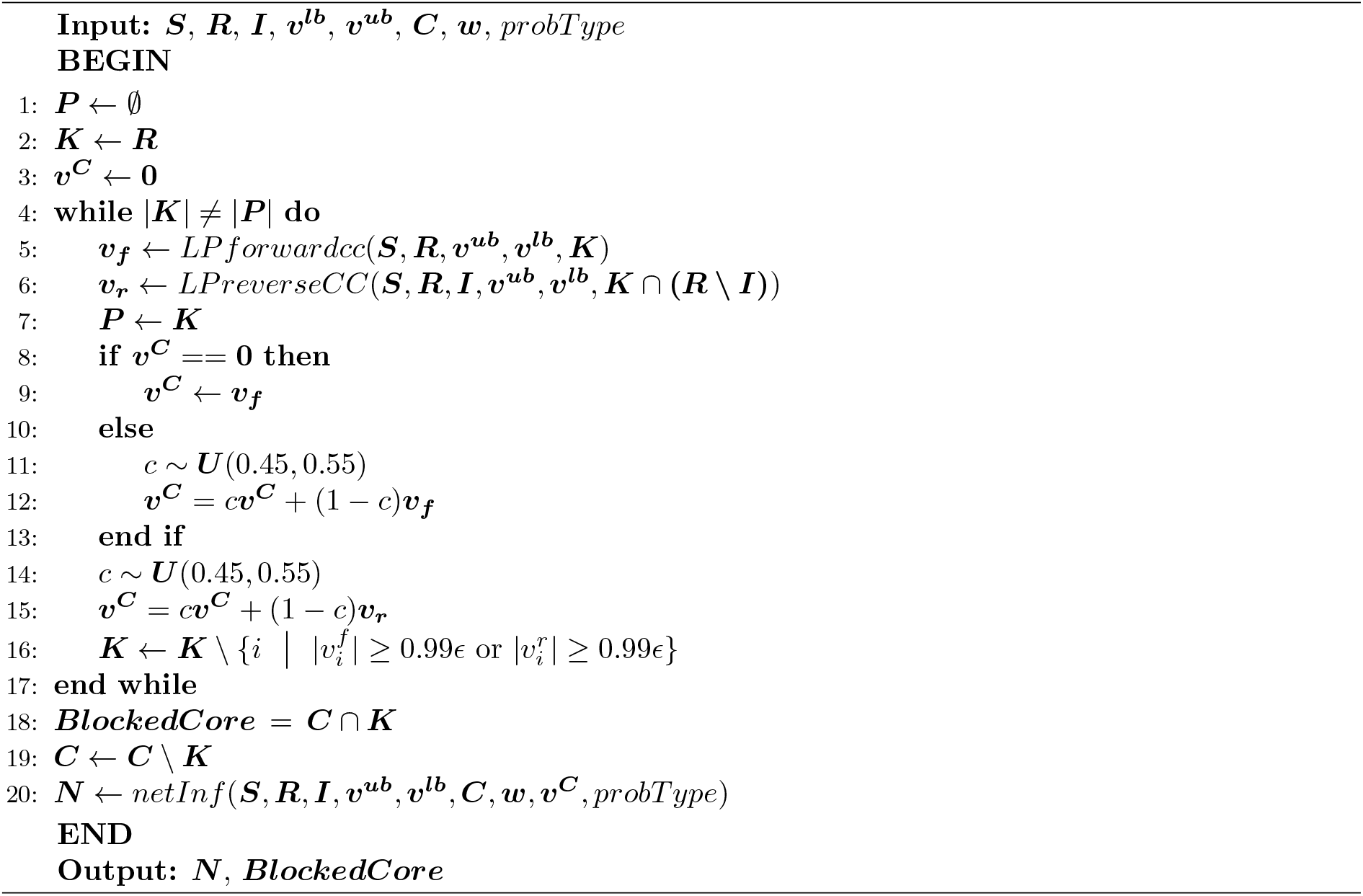

### 4.7 SPECTRA_CC: Algorithm to check the consistency of a metabolic model and report the blocked reactions

Blocked reactions typically result from dead-end metabolites or incomplete metabolic knowledge. The LP formulations, *LPforwardCC* and *LPreverseCC* can be used iteratively to detect these blocked reactions in a given metabolic network.

*LPforward* cannot be directly used to maximize the number of reactions carrying forward flux, as the presence of blocked irreversible reactions may render the LP problem infeasible. To address this,

Equation-(LPforward) is modified by removing the directionality constraints on irreversible reactions, resulting in the formulation shown in Equation-(LPforwardCC).

The pseudocode for *SPECTRA_CC* is provided in Algorithm 3. The core reaction set ***K*** is initialized as the complete reaction set ***R***. The modified forward formulation (Equation-(LPforwardCC)) and the reverse formulation (Equation-(LPreverseCC)) are executed iteratively. In each iteration, reactions with absolute flux values greater than 0.99 *× ϵ* are excluded from further consistency checks in the subsequent iteration.

To mitigate numerical instability, the threshold *ϵ* is scaled by a factor of 0.99. The algorithm terminates when no additional reactions can be verified as consistent.

### 4.8 Benchmarking protocol for the runtime comparisons

#### Environment

All runs were performed on a single cluster partition, four cores and 64 GB of memory per task, under MATLAB R2023b with the COBRA Toolbox [35] and Gurobi 11 as the solver.

#### Solver backend

FASTERCC calls the IBM CPLEX MATLAB interface (*cplexlp*) directly rather than through the COBRA Toolbox. That interface is available only up to CPLEX 12.10, whose MEX binary failed non-deterministically under MATLAB R2023b in our tests. FASTERCC’s two LP routines

##### Algorithm 3

*SPECTRA_CC*: Algorithm to report the inconsistent reactions in a metabolic model

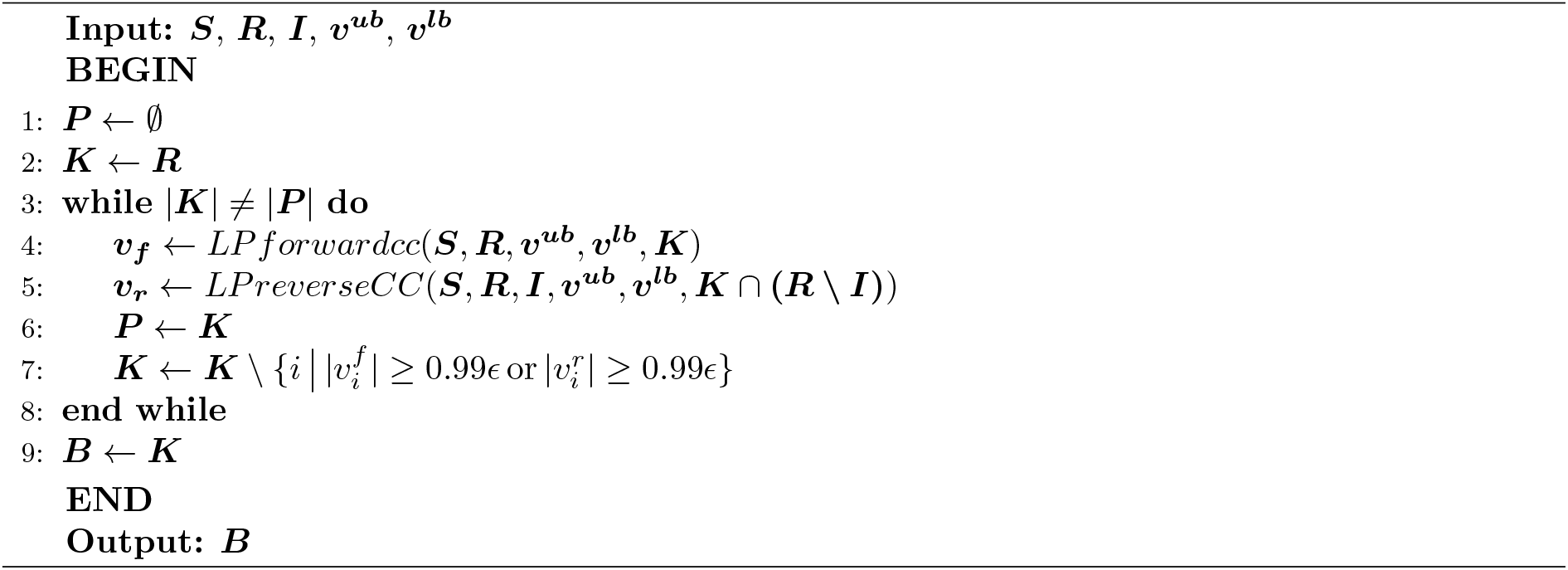

were therefore routed through *solveCobraLP* so that all methods call the same solver through the same interface. The timings reported here are consequently of FASTERCC’s algorithm on a common backend; the substitution adds the per-call overhead of the COBRA wrapper to FASTERCC and removes it from no method, so it is conservative with respect to the comparison.

#### Test sets

Three comparisons were run, all derived from Recon3D [40].

For model extraction from a consistent universal reconstruction (Fig. 2a), Recon3D was reduced to its flux-consistent core of 10,600 reactions, and core reaction sets of 20 to 10,580 reactions were sampled at random from it. SPECTRA_ME, SPECTRA_ME2, Fastcore and Swiftcore, in both its reduced and unreduced variants, were applied to each.

For model extraction from an inconsistent universal reconstruction (Fig. 2b), random universal models of 5,000 to 10,500 reactions were drawn from Recon3D, which are inherently inconsistent, and core sets of 50 to 5,000 reactions were sampled within each, giving 50 to 73 instances per universal model size. SPECTRA_CC_ME was compared against the two-step pipelines FastCC followed by Fastcore and SwiftCC followed by Swiftcore.

For consistency checking (Fig. 2c), input networks were formed from the flux-consistent core by sampling *n* reactions to retain, for *n* from 1,000 to 10,000 in steps of 1,000, and removing the remainder. SPECTRA_CC, FASTERCC, FastCC and SwiftCC were applied to each. The same construction was extended to expanded Recon3D models, generated by the scFASTCORMICS expansion for five cluster counts, and to microbial community models built by joining randomly selected AGORA2 reconstructions [44]; these are described in Supplementary Texts S1.9 and S1.10. Community members were sampled from a frozen list of all 7,302 AGORA2 models, so that an instance denotes the same community whenever it is rebuilt, and each community exchange was opened with probability 0.30 and given an inflow drawn log-uniformly between 10^*−*4^ and 10.

Every instance is a deterministic function of its size, its replicate index and a single base seed, so instances were regenerated on demand rather than stored: one expanded model at the largest cluster count exceeds 450,000 reactions.

#### Replication

For the consistency check, each condition was generated as three independent random draws and each draw timed three times, giving nine runs per condition per method; for model extraction from a consistent reconstruction, three independent core sets were drawn per size. Bars in all figures are means over those runs. SPECTRA was run without a fixed random seed, as a user would obtain it.

#### Tolerances

The detection threshold *ϵ* was 10^*−*4^ for Recon3D and expanded Recon3D models and 10^*−*8^ for community models, where the medium can limit a reaction to a flux below the higher threshold. The solver feasibility and optimality tolerances were left at their default of 10^*−*6^ for the comparisons in Fig. 2c, and set to 10^*−*9^ for the expanded and community models; the consequences of that choice are characterised in Supplementary Text S1.10. For Harvey, whose bounds span 14.3 orders of magnitude, the solver default was used, as 10^*−*9^ renders the model infeasible.

#### Adjudication

Each instance stored the reactions on which the methods disagreed, and flux variability analysis was applied to those reactions in a separate pass, under the same solver tolerance as the run being adjudicated. A reaction was taken to be consistent if FVA showed it able to carry a flux of at least *ϵ* in either direction. A method reporting a reaction consistent that FVA proves blocked is counted as an overclaim, and one reporting a reaction blocked that FVA shows carries flux as an underclaim.

### 4.9 Case Study 1: Stoichiometric consistency-based gap filling of the AGORA2 database

For both gap filling after random reaction removal and gap filling in the AGORA2 database, the same universal reconstruction was used. This universal reconstruction was defined as the union of all reactions present across the AGORA2 models. This involved three steps: (i) collecting all reactions, metabolites, and their attributes; (ii) resolving conflicts or inconsistencies in these attributes; and (iii) filtering out biomass reactions and their associated attributes.

For the first task—gap filling AGORA2 models after random deletion of 30% of their reactions, the stoi-chiometrically consistent universal reconstruction of AGORA2 was provided as input to *SPECTRA_ME*. All three minimal network–based inference algorithms were applied to this reconstruction. Let ***C***_***i***_ denote the reactions in the stoichiometrically consistent AGORA2 model *i*, ***M***_***i***_ the reactions in the corresponding random-deletion model, ***U*** the reactions in the universal reconstruction, and ***G***_***i***_ the reactions in the gap-filled model. The set of core reactions was defined as ***M***_***i***_ *∩* ***U*** . Note that reactions were removed randomly from the consistent model ***C***_***i***_ to generate ***M***_***i***_. All non-core reactions in ***U*** were assigned a weight of one, as the aim was to perform stoichiometric based gap filling across the AGORA2 database. For evaluation, we defined binary vectors representing the “actual” and “predicted” sets of recovered reactions. The vector ***actual***_***i***_ was indexed over (***C***_***i***_ *∪* ***G***_***i***_) *\* ***M***_***i***_ i.e., reactions present either in the original consistent model ***C***_***i***_ or in the gap-filled model ***G***_***i***_ but absent from the deletion model ***M***_***i***_. It was defined as:

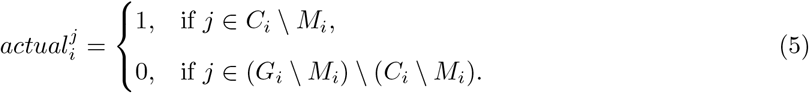

Similarly, the vector ***predicted***_***i***_ was indexed over the same set (***C***_***i***_ *∪* ***G***_***i***_) *\* ***M***_***i***_, and was defined as:

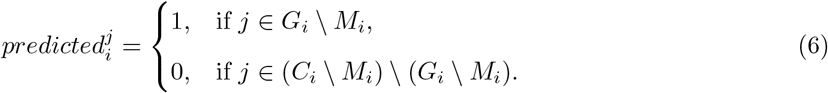

For each model *i*, we computed the standard evaluation metrics based on ***actual***_***i***_ and ***predicted***_***i***_. Let

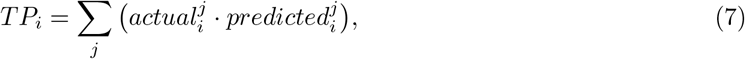

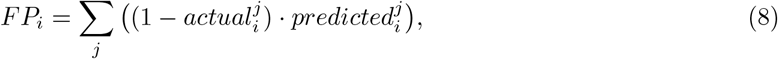

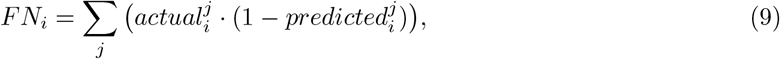

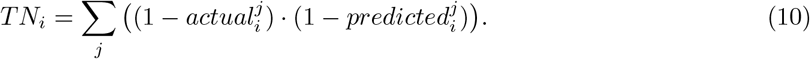

Using these, the metrics are defined as follows:

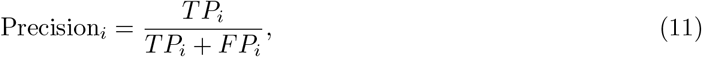

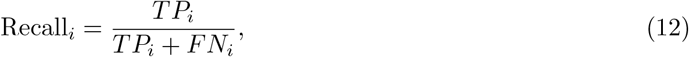

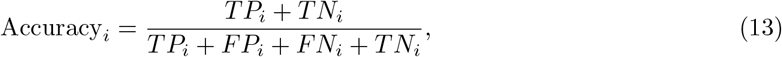

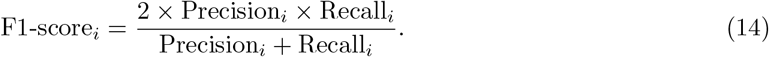

For the second task, which aims to generate AGORA2 models with fewer blocked reactions, the stoi-chiometrically consistent universal reconstruction of AGORA2 was provided as input to *SPECTRA_ME*. The core reactions were defined as all reactions from the inconsistent AGORA2 model that were also present in the universal reconstruction. Uniform weights of one were assigned to all reactions. For this task, the network inference algorithm employed was *minNetMILP* . The resulting gap-filled models were subsequently clustered by taxon based on reaction presence. To this end, binary matrices were constructed in which rows correspond to individual models and columns correspond to reactions in the universal reconstruction. Matrix entries indicate the presence (1) or absence (0) of a given reaction in a model. Two matrices were analyzed: (i) one including all reactions present in the reconstructed models, and (ii) one including only the newly added reactions. Columns containing only zeros were removed prior to further analysis. Model clustering was performed using the scikit-learn implementation of tSNE with the Euclidean distance metric.

### 4.10 ase Study 2: Microbial community gap filling

SPECTRA’s scalability at the community level was evaluated through two sets of experiments: (1) single-model and community-scale gap filling of microbes from the hCom community using CarveMe’s universal reconstructions, and (2) single-model and community-scale gap filling of hCom microbes after random reaction removal from the AGORA2 database.

#### Single-model and community gap filling using CarveMe universal models

For the first set of experiments, the protein sequences of 104 microbes were retrieved from the NCBI database. Corresponding Gram-staining information was obtained from the literature and provided in Supplementary Data S7 Data. CarveMe provides three distinct universal reconstructions for bacteria: Gram-positive, Gram-negative, and a common universal model. Based on the Gram-staining classification, the appropriate universal reconstruction was used for gap filling. All single-model and community gap-filling were performed under the standard amino acid complete (SAAC) medium, which defines the nutrient constraints on the metabolic models. The composition of the SAAC medium, including metabolite concentrations and flux bounds for both CarveMe and AGORA2 models, is detailed in Supplementary Data S8 Data. Draft genome-scale models for each microbe were generated using SPECTRA_ME using the *tradeOff* as the network inference algorithm, with the weights derived from the protein sequences and the universal reconstructions. The lower bounds of the biomass and ATP maintenance reactions were modified to have a minimal value, 0.1. This ensures the inclusion of these two reactions in the final model. No reactions were selected as a core reaction here. The reaction weights were determined based on sequence alignment scores obtained using DIAMOND [62], where each input sequence was aligned to sequences in the BiGG database [63]. Alignment scores were mapped to reactions using GPR rules and normalized to have a median of 1, following a log-normal distribution as in CarveMe. The enzymatic reactions without an expression evidence were given a score of -1, zero weights to the exchange reactions and normalized sequence alignment scores to all other reactions.

These draft models were subsequently gap filled using SPECTRA at both the single-model and community scales using *minNetLP* and *minNetMILP* as network inference methods. For single-model gap filling, no core reactions were specified, and box constraints were defined by the SAAC medium. Reaction weights were determined based on sequence alignment scores obtained in the previous step. The reaction weight *w*_*i*_ for reaction *i* was defined as:

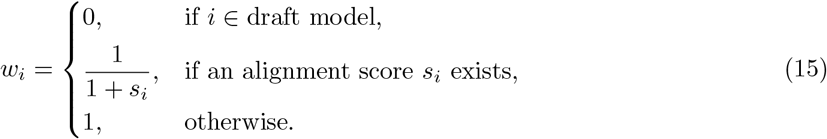

The lower bounds for the biomass and ATP maintenance reactions were set to 0.1 in all universal reconstructions. For community-scale gap filling, all exchange reactions from the respective universal reconstructions were removed, and all 104 microbial models were allowed to exchange metabolites through a shared compartment. This compartment was constrained according to the SAAC medium, and the combined model effectively represented the superset of all possible metabolic interactions within the community. The resulting community universal reconstruction contained 518,143 reactions and 235,647 metabolites. Reaction weights were assigned as defined in Equation 15. As in the single-model case, no core reactions were specified, and the biomass and ATP maintenance reactions were constrained with lower bounds of 0.1. The consistent community network extracted after gap filling was decomposed into individual organism models. For Fig. 4a, reaction counts in each model were normalized to exclude exchange reactions.

#### Community gap filling using AGORA2 models

For the second set of experiments, manually curated AGORA2 models were used to benchmark community gap filling against single-model gap filling. Of the 104 hCom species, 93 corresponding models were available in the AGORA2 database. A community model was constructed from these individual models and constrained under the SAAC medium. Stoichiometric consistency analysis was performed to obtain a consistent community model, from which the individual organism models were extracted and used as the ground truth dataset. To simulate incompleteness, 30% of reactions were randomly removed from each model. These “reaction-deletion” models were then subjected to gap filling using SPECTRA. The evaluation metrics used for performance comparison—illustrated in Fig. 4c were defined following the same framework as described in Section 4.9.

### 4.11 Case Study 3: Extraction and evaluation of cancer cell line specific models

Genome-scale metabolic models were reconstructed for 1,479 cancer cell lines using gene expression data from the Cancer Cell Line Encyclopedia (CCLE) (version: DepMap Public 23Q4). For this purpose, we used an enhanced version of the Recon3D model, referred to as Recon3D+, which includes 703 additional reactions and 25 additional genes compared to the original model [40]. The process of expanding Recon3D and curating the newly added reactions is described in Supplementary TextsS1.8. These modifications ensure that Recon3D+ maintains flux consistency while reducing the number of orphan reactions, thereby improving its suitability for context-specific model extraction. The gene–protein–reaction (GPR) rules for the newly added reactions and the complete Recon3D+ model are provided in Supplementary DataS1 Data and S2 Data, respectively.

Cancer cell line–specific models were then extracted using three stoichiometric consistency and core reactions based algorithms: Fastcore, Swiftcore, and *SPECTRA_ME*. Core reactions were defined using the Localgini thresholding algorithm [38], which assigns reactions as core based on gene expression inequality across cell lines, quantified using the Gini coefficient. In addition to identifying core reactions, Localgini also assigns weights to non-core reactions based on the expression evidence, which serve as input to the model-extraction algorithms. The performance of the algorithms was evaluated based on their ability to recover cancer hallmark reactions known to be active in cancer cell lines.

To identify cancer-specific metabolic functionalities of the extracted CSMs, we compiled a list of metabolic enzymes known to positively or negatively regulate proliferation and the Warburg effect in cancer cell lines [50]. From this curated set of experimentally validated metabolic targets, 135 reactions were successfully mapped to the Recon3D+ model and used as a benchmark to assess the coverage of cancer-specific functionality in the extracted models. These reactions are provided in the Supplementary Data S3 Data.

For further analysis, ten pairs of *SPECTRA_ME* derived models representing cancerous and normal contexts were selected. Flux Variability Analysis (FVA) was performed on each model to compute the maximum and minimum allowable fluxes for every reaction, denoted as *maxFlux*^*i*^ and *minFlux*^*i*^, respectively, for a given reaction *i*. To quantify the differential flux activity of reactions between cancerous and normal states and to further identify dysregulated reactions in cancer CSMs compared to non-cancerous CSMs, we computed the Flux Span Ratio (FSR) using the following formula:

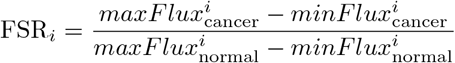

Further, to identify the dysregulated metabolic pathways, the in-built Flux Enrichment Analysis (FEA) was performed using hypergeometric 1-sided test and FDR correction for multiple testing [35]. The dysregulated reactions found using the FSR method were used to get enriched metabolic pathways in each cancer in this FEA analysis. Metabolic pathways whose adjusted *p − values* (*adj − p*) were less than 0.05 were selected as the enriched pathways. Literature analysis was carried out to get evidence of these enriched pathways. Evidences of pathways in specific cancers were categorised into ‘Pathway-level’ and ‘Component-level’ based on whether the full pathway has been implicated in the specific cancers and only a few enzymes or metabolites from the pathway were enriched, respectively. The evidence was marked ‘uncertain’ when no related information on the pathway was found in a specific cancer. A bubble plot was created by plotting cancers on the y-axis and the enriched metabolic pathways on the x-axis. The adjusted *p − values* from the FEA analysis were negative log transformed ( *−log*_10_(*adj − p*)) to denote the bubble sizes. Pathway-level, component-level and uncertain labels were given distinct colours from each other. Further, a pan-cancer metabolic analysis was carried out to describe the metabolic pathways dysregulated in more than 5 cancers.

## 5 Code availability

The codes used in this study is made open source at https://github.com/bisect-group/SPECTRA SPECTRA is distributed in two implementations. The MATLAB toolbox, which produced the results reported in this paper, is available at https://github.com/bisect-group/spectra, and the Python package at https://github.com/bisect-group/spectra-cobra. Documentation and worked tutorials for both are hosted at https://spectra-cobra-matlab.readthedocs.io/en/latest/ and https://spectra-cobra.readthedocs.io/en/lates_respectively.

## 6 Data availability

The data required to reproduce all the results present in the paper are made open source at https://github.com/bisect-group/SPECTRA.

The gap-filled AGORA models with lesser blocked reactions can be obtained from https://doi.org/10.5281/zenodo.18126152

## 7 Supplementary information

**S1 Supplementary information** Supplementary information containing supplementary text, supplementary tables, and supplementary figures.

**S1 Data** Gene protein reactions rules for the newly appended reactions in Recon3D+ model.

**S2 Data** Recon3D+ model in .mat format.

**S3 Data** The 135 cancer core reactions that served as a reference set to evaluate the performance of the consistency-based model extraction methods.

**S4 Data** The list of reactions that are up-regulated in the cancer-cell lines, along with their Flux Span Ratio (FSR) values and the associated gene names.

**S5 Data** The list of reactions that are down-regulated in the cancer-cell lines, along with their Flux Span Ratio (FSR) values and the associated gene names.

**S6 Data** The list of subsystems that are significantly deregulated in the cancerous cell lines compared to the normal cells.

**S7 Data** The gram staining information of 104 microbes in the hCom microbial community

**S8 Data** The molecular components with their respective concentrations and the bounds used for CarveMe and AGORA2 models.

**S9 Data** The details on number of newly added reactions and the count of reactions with EC number evidence from the CLEAN model predictions.

## 8 Acknowledgments

We gratefully acknowledge the Centre for Integrative Biology and Systems Medicine (IBSE) at IIT Madras for providing access to high-performance computing facilities essential for this work. We also thank Mitacs and the IIT Madras for their financial support to support the travel of SPK. RM and NA also acknowledge funding from Canada Research Chairs Program, NSERC and Genome Canada. The benchmarking simulations reported in this work were performed on the Digital Research Alliance of Canada (formerly Compute Canada) advanced research computing infrastructure. This project is partially supported by the University of Toronto’s Eric and Wendy Schmidt AI in Science Postdoctoral Fellowship, a program of Schmidt Sciences.

