## Supplementary information for "SPECTRA: Generalist framework for reconstruction of high-quality metabolic networks at large-scale from multi-omics data"

**S2 Supporting Tables** **13**

**S3 Supporting Figures** **23**

### S1 Supplementary text

#### S1.1 Rationale behind using random numbers in the objectives of LPforward, LPforwardCC and LPreverseCC

To explain the rationale behind using the random numbers in the objective of the LPforward, LPforwardCC and LPreverseCC, we use a toy model and an LP formulation, LPforward_0 without the random weights. For a given input of reactions $\boldsymbol{K}$ ($\subseteq \boldsymbol{R}$), Equation-(LPforward_0) shown below can maximize the number of reactions that can carry a positive flux. In certain conditions LPforward_0 cannot bring up positive fluxes through the reaction. One such example is shown in Figure S1.

$$
\begin{aligned}
\max_{\boldsymbol{v},\boldsymbol{z}} \quad & \sum_{i \in \boldsymbol{K}} z_i \\
\text{s.t.} \quad & \boldsymbol{S}\boldsymbol{v} = \boldsymbol{0} \\
& v_i^{lb} \leqslant v_i \leqslant v_i^{ub} \quad && \forall i \in \boldsymbol{R} \\
& v_i \geqslant z_i && \forall i \in \boldsymbol{K} \\
& z_i \leqslant \epsilon && \forall i \in \boldsymbol{K}
\end{aligned}
\tag{LPforward_0}
$$

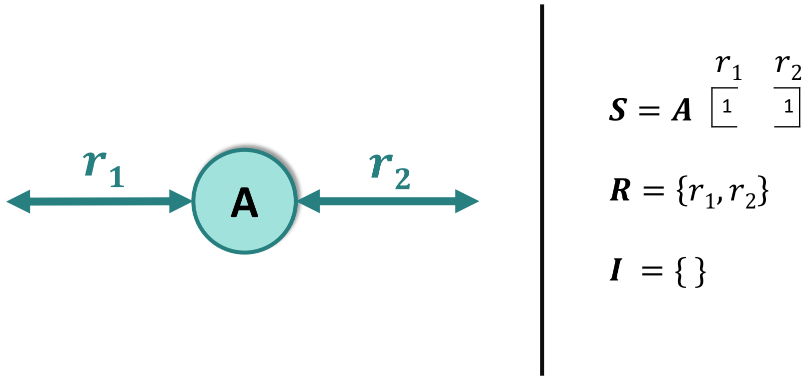

Figure S1: **Example metabolic network where Equation-(LPforward_0) fails to maximize the flux through the reactions**

Here, if both the reactions $r_1$ and $r_2$ belong to $\boldsymbol{K}$ then solution to the objective of LPforward_0 is zero always. This could result in solution being a zero vector. The objective function is modified to account for this zero vector solution. In LPforward the decision variables $z_i$ are given random weights $x_i$. For all the simulations in this study, the random weights are chosen from the uniform distribution $\sim \boldsymbol{U}(1, 1.1)$. Also, note that in this formulation, all the irreversible reactions are forced to carry a flux in forward direction. This is because, a single run of LPforward can make all the irreversible reactions have a positive flux.

$$\begin{aligned}
\max_{\boldsymbol{v},\boldsymbol{z}} \quad & \sum_{i \in \boldsymbol{K} \text{ and } i \notin \boldsymbol{I}} x_i . z_i \\
\text{s.t.} \quad & \boldsymbol{S}\boldsymbol{v} = \boldsymbol{0} \\
& v_i^{lb} \leqslant v_i \leqslant v_i^{ub} && \forall\, i \in \boldsymbol{R} \\
& v_i \geqslant z_i && \forall\, i \in \boldsymbol{K} \text{and}\, i \notin \boldsymbol{I} \\
& z_i \leqslant \epsilon && \forall\, i \in \boldsymbol{K} \text{and}\, i \notin \boldsymbol{I} \\
& v_i \geqslant \epsilon && \forall\, i \in \boldsymbol{K} \text{and}\, i \in \boldsymbol{I}
\end{aligned} \tag{LPforward}$$

#### S1.2 LPforwardCC in SPECTRA is faster compared to LP7 in Fastcore

At the core of Fastcore for model reconstruction is the linear program, Equation-(LP7). Equation-(LP7) detects the unblocked reactions that can carry a positive flux. Fastcore iteratively solves LP7 on a given input model. Building upon LP7, LPforwardCC was developed by adjusting constraints on the decision variable $z_i$ and introducing random weights within the objective function. Both LP7 and LPforwardCC are assessed based on number of unblocked reactions identified in a single run.

$$\begin{aligned}
\max_{\boldsymbol{v},\boldsymbol{z}} \quad & \sum_{i \in \boldsymbol{K}} x_i . z_i \\
\text{s.t.} \quad & \boldsymbol{S}\boldsymbol{v} = \boldsymbol{0} \\
& v_i^{lb} \leqslant v_i \leqslant v_i^{ub} && \forall\, i \in \boldsymbol{R} \\
& v_i \geqslant z_i && \forall\, i \in \boldsymbol{K} \\
& z_i \leqslant \epsilon && \forall\, i \in \boldsymbol{K}
\end{aligned} \tag{LPforwardCC}$$

Subnetworks derived from the Recon3D model [1], varying in size from 20 to 10600, were given as input to LP7 and LPforwardCC. Each LP was solved once, and the number of identified unblocked reactions by both LPs is depicted in Figure S2. The results illustrate that LPforwardCC outperforms LP7 by identifying a higher number of unblocked reactions. Furthermore, the LP formulations in SPECTRA improves pathway coverage by activating most reactions within individual pathways, as detailed next.

$$\begin{aligned}
\max_{\boldsymbol{v},\boldsymbol{z}} \quad & \sum_{i \in \boldsymbol{K}} z_i \\
\text{s.t.} \quad & \boldsymbol{S}\boldsymbol{v} = \boldsymbol{0} \\
& v_i^{lb} \leqslant v_i \leqslant v_i^{ub} && \forall\, i \in \boldsymbol{R} \\
& v_i \geqslant z_i && \forall\, i \in \boldsymbol{K} \\
& 0 \leqslant z_i \leqslant \epsilon && \forall\, i \in \boldsymbol{K}
\end{aligned} \tag{LP7}$$

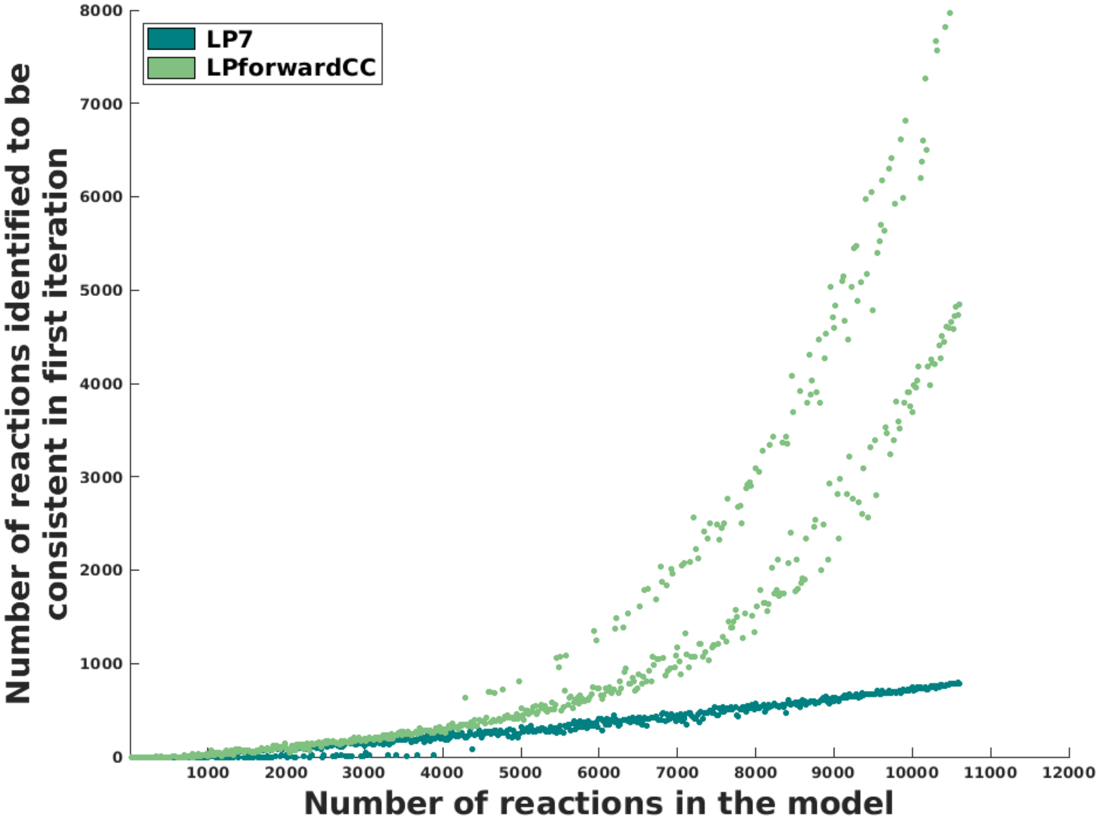

Figure S2: Number of unblocked reactions identified by solving LPforward is more than that obtained by solving LP7 for a single iteration

#### S1.3 Comparison of pathway coverage between Fastcore and SPECTRA on a toy metabolic network

Fastcore iteratively alternates between unblocking core reactions and minimizing the universal reconstruction until all core reactions are made flux consistent. However, building on LP7, Fastcore's inherent limitation in unblocking less number of core reactions in a single iteration (see section S1.2) often causes it to focus on sub-pathways rather than complete pathways. SPECTRA overcomes this through two key strategies: (1) using random-weighted objectives to unblock most core reactions simultaneously, and (2) performing network minimization only once after all core reactions are unblocked.

This difference is illustrated using the toy model in Figure S3, which contains nine reactions and five metabolites. For the same core reactions ($R_2$, $R_4$, $R_5$), Fastcore and SPECTRA (minNetLP) produce different subnetworks. Fastcore unblocks and minimizes one pathway segment at a time, yielding fragmented pathways. In contrast, SPECTRA first unblocks all core reactions using the combined information from LPforwardCC (or LPforward) and LPreverse (see Methods), then performs network inference optimization. This comprehensive unblocking strategy enables SPECTRA to capture complete pathway information before network minimization, resulting in more biologically coherent reconstructions.

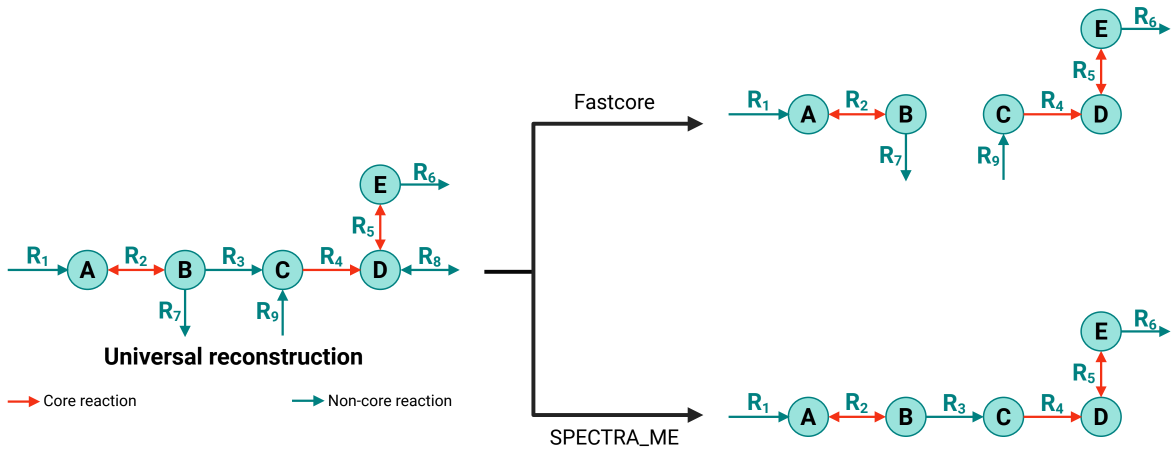

Figure S3: The universal reconstruction is a toy model with nine reactions ($R_1$–$R_9$) and five metabolites, with core reactions $R_2$, $R_4$, $R_5$ indicated. Fastcore's iterative unblocking and minimization produces fragmented sub-pathways. On the otherhand, SPECTRA first comprehensively unblocks all core reactions then performs single-step network minimization (minNetLP), capturing the complete pathway structure.

#### S1.4 Swiftcore does not consider constraints on non-core reversible reactions

In a simplified genome-scale metabolic model (GEM) containing seven reactions and five metabolites as depicted in Figure S4, constraints other than thermodynamics are considered. These constraints include i) Exchange reaction constraints (applied to reactions $R_1$ and $R_4$), representing environmental and media conditions, and ii) Constraints on an irreversible reaction (applied to reaction $R_6$), which might reflect constraint on ATP-maintenance reaction, biomass reaction, or other regulatory constraints. The core reaction in this case is assumed to be $R_6$.
The substrate, $\boldsymbol{A}$ involved in reaction $R_6$ can be produced through two distinct pathways: one pathway comprises reactions $R_1$, $R_2$, and $R_3$, while the other involves reactions $R_4$ and $R_5$. For the core reaction $R_6$ to exhibit flux within the provided bounds, both pathways must remain active.
Both SPECTRA and Fastcore consider all bounds specified within the GEM. Consequently, these algorithms construct a context-specific model with both the pathways. However, Swiftcore, by disregarding constraints on non-core reversible reactions, specifically ignores constraints associated with media conditions. As a result, Swiftcore might generate a minimal model comprising only one pathway (reactions $R_4$ and $R_5$). This would render an infeasible model due to the lower bounds imposed on reactions $R_4$ and $R_6$.
Indeed, exchange reactions includes both active and passive transport reactions. Given that passive transport reactions have no genes or proteins to have expression evidence, they are not considered in the list of core reactions. In scenarios like these, Swiftcore might generate an infeasible model due to the exclusion of constraints related to media conditions.

#### S1.5 SPECTRA - a flexible framework for network inference: Application to a Toy Network

The core of SPECTRA consists of eight optimization formulations that provide flexibility in both constraints and objectives. Three LP formulations: `LPforward`, `LPforwardCC`, and `LPreverseCC` are

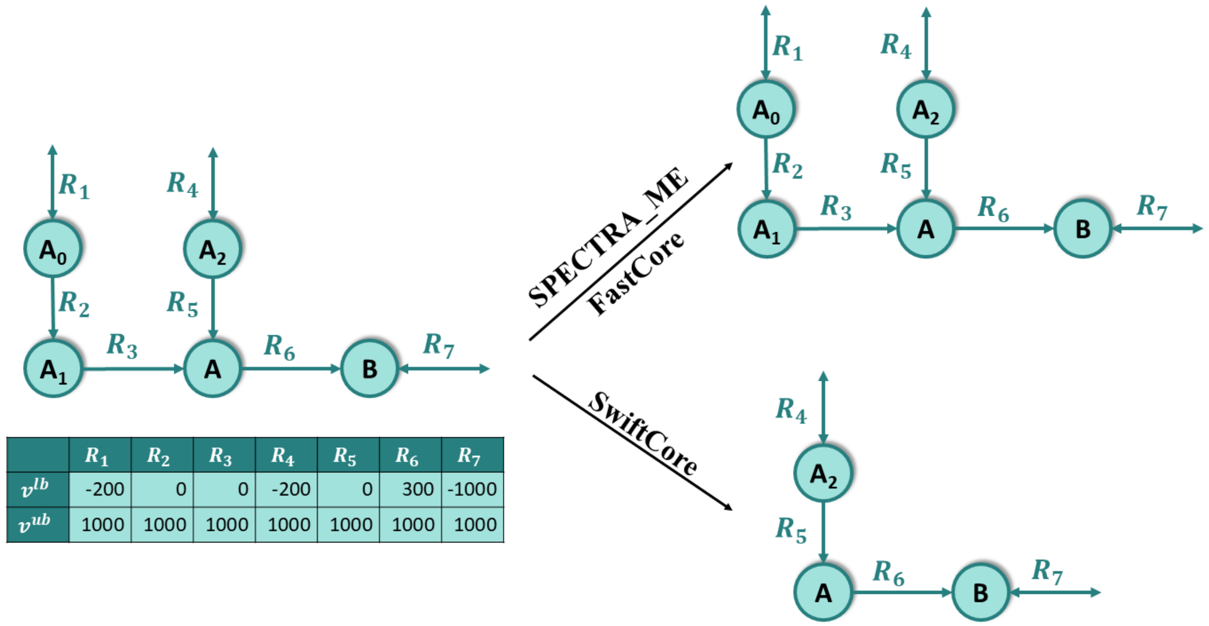

| | $R_1$ | $R_2$ | $R_3$ | $R_4$ | $R_5$ | $R_6$ | $R_7$ |
|---|---|---|---|---|---|---|---|
| $v^{lb}$ | -200 | 0 | 0 | -200 | 0 | 300 | -1000 |
| $v^{ub}$ | 1000 | 1000 | 1000 | 1000 | 1000 | 1000 | 1000 |

Figure S4: An illustration of a metabolic network for which Swiftcore builts an infeasible context-specific model

iteratively applied to obtain a feasible flux vector that enforces nonzero flux through the specified core reactions. This flux vector is then used as a constraint in five distinct network inference formulations. To further illustrate the method and the utility of these formulations, we use a toy network shown in Fig. S5. In the toy example, the network contains eight metabolites and eleven reactions. The secretion of metabolite D (reaction $R_5$) is designated as a core reaction, which can be achieved via three alternative pathways: P1 ($R_1$–$R_2$–$R_3$–$R_4$), P2 ($R_6$–$R_7$–$R_8$), and P3 ($R_9$–$R_{10}$–$R_{11}$).

For the LP-based formulation minNetLP, the inferred subnetwork is pathway P1, despite involving more reactions than P2 or P3. This outcome arises from stoichiometric considerations: reactions $R_8$ and $R_{11}$ generate only 0.5 mol of metabolite D, requiring higher flux levels in P2 and P3 compared to P1. Reaction weights also influence pathway selection by acting as penalties—higher weights reduce the likelihood of a reaction being included. Importantly, for minNetLP, all weights must be non-negative; otherwise, the formulation becomes unbounded.

In contrast, the MILP formulation minNetMILP selects subnetworks based solely on the number of reactions. Thus, P2 or P3 both with fewer reactions are favored over P1. As with minNetLP, weights act as penalties and must be non-negative. MILP-based methods such as minNetMILP and tradeOff can also generate alternative optimal (or sub-optimal) solutions by iteratively excluding at least one reaction from prior solutions. In the toy network, the first solution is P2 (four reactions), the second is P3 (also four reactions, equally optimal), and the third is P1 (five reactions, suboptimal). The minNetDC formulation addresses the problem as a cardinality optimization and is solved using the difference-of-convex approach described in [2].

The tradeOff MILP formulation instead optimizes the total weight of the selected pathway. With reaction weights assigned as $[R_1–R_{11}] = [1, 1, -2, -1, 1, 1, 2, -1, -2, 0, 1]$, P1 achieves a total weight of –1, P2 achieves 2, and P3 achieves –1. The algorithm therefore returns subnetworks in decreasing order of total weight: P2 first, followed by P1 or P3. In this case, weights can be any real value, including negatives. If all pathway weights are positive, the complete universal model will be returned in the first solution.

Finally, optimBiomass, the LP formulation balances the inclusion of reactions with maximizing biomass flux. Reaction weights here serve dual purposes: penalizing reaction inclusion (as in minNetLP) while also determining the tradeoff with biomass production. For example, assigning a high weight (10) to the biomass reaction ($R_5$) results in inclusion of all reactions, since they collectively contribute flux to biomass. Reducing the biomass weight to 1 instead favors pathway P1. In this formulation, weights must be non-negative; higher biomass weights increase biomass flux, while higher weights on other reactions reduce their likelihood of inclusion.

While alternate solutions shown here are only for MILP formulations (Pathway exclusion), we have devised a method to infer alternative solution for LP formulations (Core reaction directionality). Explanation of the method with a toy network is shown in Supplementary section S1.6. This toy example demonstrates SPECTRA's flexibility in extracting biologically meaningful metabolic subnetworks under different optimization criteria.

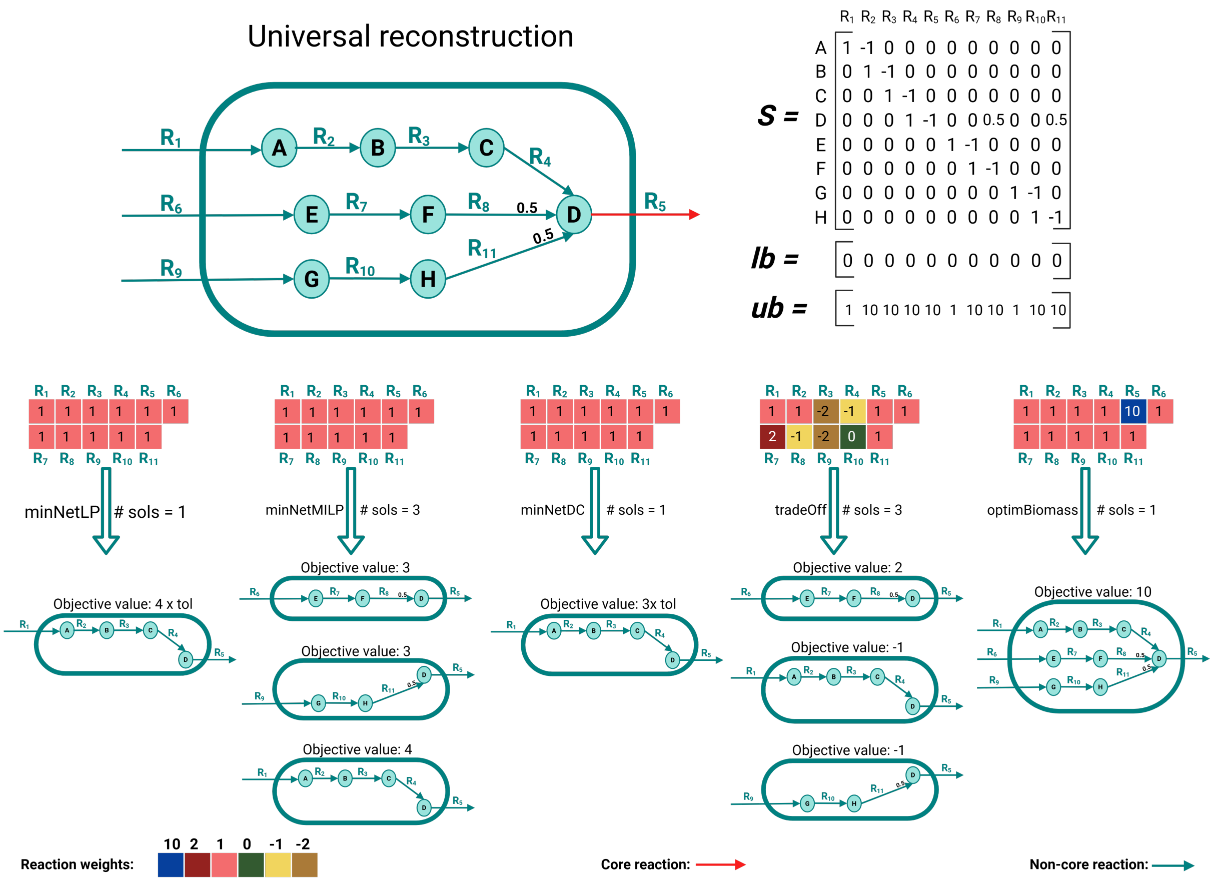

Figure S5: **SPECTRA's network inference formulations implied on the toy network** SPECTRA provides five optimization formulations each with distinct objective for the network inference. The toy network shows three pathways that can produce the metabolite D (similar to the biomass precursor). Based on the provided weights (colored squares) and the core reaction details (Here reaction $R_5$ is core reaction) and the bounds on the universal reconstructions, each of the formulations infer distinct pathways. Here, # sols refer to the number of alternate solutions required.

#### S1.6 Algorithm to get alternative solutions for a given set of core reactions:

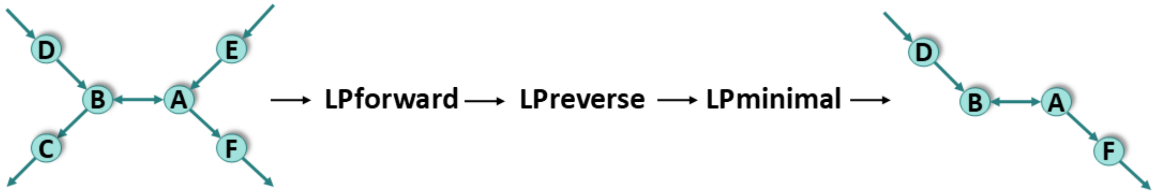

(a) The context-specific model is constructed by first solving LPforward followed by LPreverseCC.

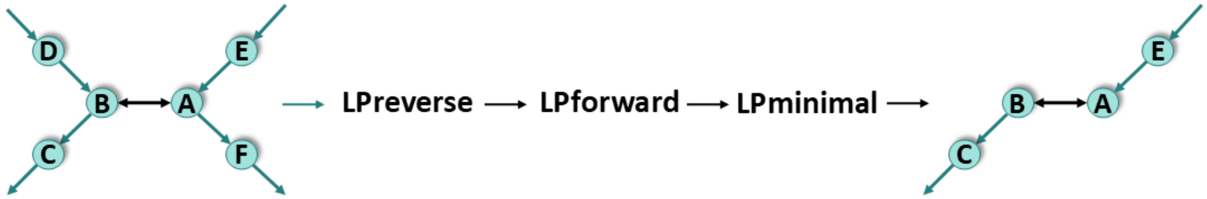

(b) The context-specific model is constructed by first solving LPreverseCC followed by LPforward.

Figure S6: The order of solving LPforward and LPreverseCC impacts the resulting context-specific models, generating differences between them

For a given set of core reactions, SPECTRA_ME (or SPECTRA_CC_ME) can generate different optimal subnetworks when multiple solutions exist for LPforward, LPreverseCC, or LPforwardCC, effectively making the algorithm stochastic. Additional alternative subnetworks can also be obtained by changing the order in which LPreverseCC and LPforward (or LPforwardCC) are solved (Fig. S6). This reordering yields different flux vectors $\boldsymbol{v}^{\boldsymbol{C}}$ (see Methods section), where the flux directions of core reversible reactions may vary.

To systematically capture such alternative minimal networks, we developed a formulation that allows users to request multiple solutions. In this approach, the order of solving LPforward and LPreverseCC is randomized in each iteration, thereby generating distinct $\boldsymbol{v}^{\boldsymbol{C}}$ vectors. It is important to note, however, that this strategy does not apply when all core reactions are irreversible, since in such cases the solution is unique.

#### S1.7 Kinds of blocked reactions

Blocked (or inconsistent) reactions can arise either from the structure of the network or from imposed flux bounds. Structural causes are of two kinds: stoichiometric and topological. Stoichiometric constraints enforce the pseudo–steady-state assumption, ensuring that metabolites do not accumulate indefinitely. Topological constraints, in contrast, require that reactions remain connected within the network. For example, in the first toy network shown in Fig. S7 (six metabolites and six reactions), the stoichiometric coefficient of the reaction $\mathsf{S} \rightarrow \mathsf{n}\mathsf{A}+\mathsf{B}$ determines whether the network is consistent. When $\mathsf{n} = 1$ or $\mathsf{n} = 3$, the stoichiometry constraints results in all reactions becoming blocked. Under topological constraints, however, the network remains consistent regardless of the value of $\mathsf{n}$, since metabolite accumulation is permitted: metabolite $\mathsf{B}$ can accumulate when $\mathsf{n} = 1$, and metabolite $\mathsf{A}$ when $\mathsf{n} = 3$. However, neither stoichiometric nor topological constraints allow unrestricted consumption of metabolites. This

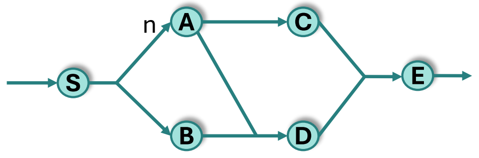

| | Stoichiometry | Topology |
|---|---|---|
| n=1 | Inconsistent | Consistent |
| n=2 | Consistent | Consistent |
| n=3 | Inconsistent | Consistent |

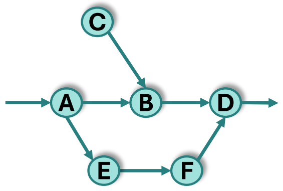

| Stoichiometry | Topology |
|---|---|
| Inconsistent | Inconsistent |

Figure S7: Toy networks describing the kinds of blocked reactions. In the first toy network, the choice of the stoichiometric coefficient, $n$ determines the consistency of the network if stoichiometric constraints are imposed and this is not the case of topology based constraints. Both the topology constraints and the stoichiometric constraints do not allow for free consumption of the metabolites as shown in the second toy network. The values in the tables indicate the consistency of the entire network

is illustrated in the second toy network (six metabolites, eight reactions), where the reaction $C \rightarrow B$ is blocked because metabolite $C$ has no producing reaction. Blocked reactions can also arise from flux bounds, which are condition-specific rather than structural. For example, in the second toy network, if the import of metabolite $A$ is constrained to zero flux, then the entire network becomes inconsistent.

SPECTRA_CC enables identification of all such types of blocked reactions by explicitly defining both the reaction bounds and the type of structural constraint to be imposed.

#### S1.8 Improved reconstruction of the Recon3D model for constraint-based applications

Genome-scale metabolic models (GEMs) of human tissues and cell lines are typically reconstructed by integrating transcriptomic data into whole body consensus metabolic networks using various algorithms. These consensus networks such as Recon3D [1] and Human GEM [3] comprise extensively curated reactions and gene associations collected from literature. Compared to manual curation, which is time-consuming and labor-intensive, the use of omics data to generate context-specific models (CSMs) from these consensus reconstructions has become standard practice.

The Virtual Metabolic Human (VMH) database hosts two versions of the Recon3D model [4]: the Recon3D reconstruction (Recon3D_full) and the Recon3D model (Recon3D_consistent). While the consistent model is suitable for constraint-based modeling, the full reconstruction contains issues such as blocked reactions, flux inconsistency, and violations of sanity checks—tests that ensure the model does not have erroneous energy reactions that leak protons, cause excess ATP, etc [1, 2].

However, the consistent Recon3D model is significantly smaller than the full reconstruction; it includes 3000 fewer reactions, leaving room to augment its size without compromising model consistency. To this end, we developed an improved version of the curated Recon3D model, referred to as Recon3D+, by incorporating additional reactions from the full reconstruction. Below, we describe the steps taken to enhance the model:

1. Removal of duplicate reactions: Prior to adding reactions from the Recon3D_full to Recon3D_consistent version, duplicate reactions were removed from both the models. These included reactions with identical stoichiometry but differing in directionality (reversible vs. irreversible) and transport reactions where substrates and products were reversed. Using the COBRA Toolbox [5] functions, we identified and removed 243 such reactions. The retained counterpart transport reactions were all made reversible to enable unconstrained inter-compartmental transport of the metabolites.
2. Pruning of unused genes: Genes not associated with any reactions were removed from both models.
3. Addition of reactions: Reactions from the Recon3D_full reconstruction were added into the Recon3D_consistent model iteratively, ensuring that each addition passed standard sanity tests [5]. These tests verify that the model does not exhibit unrealistic energy production (e.g., generating ATP from water or without substrate input). An iterative validation process was employed to identify and filter out problematic reactions.
4. Elimination of redundant and erroneous reactions: Specific reaction sets were removed to improve model fidelity. These included unbounded NAD(P)H quinone dehydrogenase 1 (*NQO1*) futile cycle reactions—such as NADPQNOXR, NADQNOXR, HMR_9538, and HMR_9722 which resulted in incorrect NADPH generation, and oxalosuccinic acid–producing IDH1 reactions (r0422–r0425), which overlapped with ketoglutarate-producing variants [6].
5. Flux consistency refinement: Finally, stoichiometrically inconsistent reactions were removed after the completion of the previous steps, to ensure that the resulting model (Recon3D+) was flux-consistent and suitable for use with consistency-based algorithms such as SPECTRA_ME, Fastcore and Swiftcore.

##### S1.8.1 Updating Gene-Protein-Reaction (GPR) associations for orphan reactions in the Recon3D+ Model

A key aspect of reconstructing context-specific genome-scale models lies in accurately mapping genes to their associated reactions through gene-protein-reaction (GPR) rules. Most omics integration algorithms select reactions associated with highly expressed genes while pruning those linked to lowly expressed genes [7, 8, 9]. Therefore, reducing the number of orphan reactions; by updating gene associations to them improves the specificity and biological relevance of the resulting CSMs.

To address this, several strategies were employed to assign GPR rules to orphan reactions in the Recon3D+ model:

1. **MetaCyc and Related Databases** [10, 11, 12] MetaCyc is a comprehensive database of curated metabolic pathways and reactions, including detailed information on genes, enzymes, and transporters. Its human-specific subset, HumanCyc, provides gene-reaction associations for Homo sapiens [12]. Out of 3973 orphan reactions in Recon3D, 1546 were found to have enzyme commission (EC) numbers. Upon cross-referencing these EC numbers with other biochemical databases—including KEGG, BRENDA, BioCyc, BIGG, RHEA, and MetaNetX [13, 14, 15, 16, 17, 18] many were found to be incorrectly assigned. The first step was to rectify these EC numbers and then retrieve the associated genes from HumanCyc. The reaction IDs of Recon3D were first converted to respective database IDs, and then the EC numbers were extracted for these reactions. Then, gene annotations from HumanCyc, originally in official gene symbol format, were mapped to the Recon3D gene ID format. When multiple genes were associated with a reaction, they were combined using an OR relationship in the GPR rule.

2. **Harvetta Model** [19] Some orphan reactions in Recon3D were updated with GPR rules in the Harvetta model (sex-specific whole body metabolic reconstruction). By comparing reaction names, nine of these updated GPRs were transferred to Recon3D+.

3. **Human-GEM Model** [3] While Recon3D and Human-GEM share many reactions and metabolites, direct comparison is hindered by differences in nomenclature. However, early versions of both models contain reaction identifiers prefixed with "HMR_", allowing limited cross-model mapping. For a subset of orphan reactions in Recon3D+ with HMR_ identifiers, GPR information was available in Human-GEM. These GPR rules—originally annotated with Ensembl gene IDs—were converted to the Recon3D gene ID format and updated into the model. In total, 48 such reactions were updated with GPR rules.

In conclusion, genes for 303 orphan reactions were updated, and thereby 25 additional genes were added to Recon3D+. The gene–protein–reaction (GPR) rules for the newly added reactions and the complete Recon3D+ model are provided in Supplementary Data.

#### S1.9 SPECTRA_CC scales to half a million reactions

The consistency check in main text operates on models of the size of a single genome-scale reconstruction. Multicellular applications like microbial communities, multi-tissue networks and whole-body models are one to two orders of magnitude larger, and it is at that scale that the cost of consistency checking determines whether a reconstruction pipeline is usable at all. We therefore extended the comparison to expanded Recon3D models generated by the scFASTCORMICS expansion [20]. Each such model represents a multicellular system as $k$ copies of the base Recon3D network, one per cell cluster, with the copies connected to one another by inter-cluster transport reactions and sharing a single set of exchange reactions with the environment. The cluster count $k$ therefore sets how many cell types the model contains, and because the shared transport and exchange reactions do not scale with it, the reaction count is affine in $k$: $n(k) = 9040k + 3120$. Five cluster counts giving 47,820 to 454,620 reactions were used; in each, 500, 2,000, 5,000 or 10,000 reactions were deleted at random to plant known blocked reactions, with three independent draws per condition and three timings per draw.

Only SPECTRA_CC and FASTERCC were carried into this comparison. FastCC was excluded on runtime: it already required 3,824 s in total at Recon3D scale against 14.8 s for SPECTRA_CC, and its LP count grows with the size of the network, so at half a million reactions it is not practical. SwiftCC was excluded because it does not solve the same problem: it enforces only thermodynamic (irreversibility) constraints and does not account for bounds on exchange reactions or on reactions with positive lower bounds, so it reports as consistent reactions that those constraints block (Supplementary Table S1). It consequently returns a larger consistent set than the other methods, and a runtime comparison against it at this scale would not be comparing like with like.

SPECTRA_CC solved exactly 10 LPs at every size from 93,020 reactions upward (Supplementary Fig. S8). The count did not change across a 4.9-fold increase in model size, nor across a 20-fold change in the number of reactions deleted, and the consistent sets returned by the three timings of each draw were identical. FASTERCC solved 75–79 LPs at the two sizes it completed and returned no result at 183,420 reactions or above. The cause is not its consistency algorithm, which completed on the 183,420-reaction model with a resident set of 3.23 GB, but a step that follows it: *restoreBounds* evaluates `all(S_orig == S_cons, 1)`, and in MATLAB a comparison between two sparse matrices is true wherever both operands are zero, so the result carries one entry per matrix *element* rather than one per nonzero. Measured directly, its peak memory is approximately 7 bytes per element and independent of the number of nonzeros, predicting 29 GB at 93,020 reactions, which completes, and 112 GB at 183,420, which does not, against a 64 GB limit. At 454,620 reactions MATLAB reports the array it declines to allocate, for example 216,093 × 453,736 (821.8 GB), dimensions that are exactly the metabolite and reaction counts of the consistent model. The cost therefore scales with the size of the consistent set rather than of the input, which is limiting here because these models are approximately 99% consistent.

#### S1.10 Effect of the solver feasibility tolerance

Consistency checking decides whether a reaction can carry a flux of at least $0.99\epsilon$, so its answer depends on the feasibility tolerance of the underlying solver. We characterised this dependence on microbial community models, where it matters most. Each community model was built by joining a number of randomly selected AGORA2 reconstructions [21] into a single stoichiometric model: every organism retains its own intracellular compartments, all organisms share one common extracellular compartment through which they exchange metabolites with each other, and community-level exchange reactions connect that shared compartment to the environment. A random medium was then imposed on those community exchanges, so that reactions become blocked through the nutritional environment rather than through deletion. The smallest inflow a community exchange may be given is $10^{-4}$, which is close to the feasibility tolerances under test; this proximity is what makes the looser settings unreliable here, since a reaction limited by the medium carries a flux of the same order as the tolerance that decides whether it carries one at all. Communities of 50 and 100 organisms were used, giving 79,948 to 180,874 reactions, with three independent draws and three timings at each of four tolerances, and SPECTRA_CC run alone (Supplementary Table S2).

Only at $10^{-9}$ is the answer reproducible. Across three repeated runs of the same model the consistent set varied by at most 45 reactions at 50 species and 129 at 100 species, and in several draws was identical. At every looser tolerance the same model returned materially different answers on repeated runs, by up to 5,097 reactions at 50 species and 19,664 at 100 species. The looser settings are also more expensive, not less: SPECTRA_CC required 10-13 LPs at $10^{-9}$ and 155-476 at the other three, with runtimes 11- to 24-fold higher. The dependence is not monotone, $10^{-8}$ returns more consistent reactions than $10^{-9}$ and $10^{-7}$ fewer, which is consistent with solver residuals being read as flux once the detection cutoff approaches the feasibility tolerance.

Tightening the tolerance is therefore the correct choice for SPECTRA_CC wherever the model is well scaled, as it costs nothing in LPs and is the only setting at which the result is stable. The qualification matters: on models that are not scaled well, such as Harvey (Supplementary Text S1.11), a tolerance of $10^{-9}$ renders even flux balance analysis infeasible, and the solver default must be used instead.

#### S1.11 Consistency checking on models with forced fluxes

A flux-consistent subnetwork is by definition entirely flux consistent, so a consistency checker applied to its own output must return every reaction it is given. We implied this test on the whole-body reconstruction Harvey [19], running each method, rebuilding the model from the reactions it reported consistent, and running it again until the answer stopped changing (Supplementary Table S3).

Harvey comprises 81,094 reactions, 56,452 metabolites and 2,071 genes, of which 31,712 reactions are reversible and 1,539 are fixed to a single flux value. Its bounds span 14.3 orders of magnitude, from $5.5 \times 10^{-9}$ to $10^{6}$ in absolute value, which is why the solver tolerance cannot be tightened on it. Most importantly for this test, 167 of its reactions have a positive lower bound (23) or a negative upper bound (144) and so exclude the zero flux vector: 88 diet uptake reactions, 49 renal blood exchanges, 12 blood-brain barrier transports, water, carbon dioxide and urea excretion, cardiac ATP maintenance, and the whole-body objective, which is fixed at 1. It is the only model used in this work with that property.

Neither method was idempotent on the first pass: SPECTRA_CC reported 124 reactions as consistent that its own second pass removed, and FASTERCC 147. Both converged on the second pass. Applied to the model as distributed, however, SPECTRA_CC's second pass returned no consistent reactions at all. This is a property of the forced fluxes rather than of the method: deleting the reactions found blocked in the first pass removes carriers that the positive lower bounds require, so the trimmed network admits no feasible flux distribution. Relaxing those bounds so that the zero vector is feasible, as FASTERCC does internally to its own input restores convergence. The consistent subnetwork of a model with forced fluxes is therefore not itself a feasible model, which should be accounted for when such a subnetwork is used as the basis of a reconstruction.

### S2 Supporting Tables

Table S1: FVA adjudication of disagreements between the four consistency checking methods on random subsets of Recon3D, at two solver feasibility tolerances. Overclaim: the method reports a reaction as consistent that FVA proves blocked. Underclaim: the method reports a reaction as blocked that FVA shows can carry flux.

| | **feasTol $10^{-6}$** | | **feasTol $10^{-9}$** | |
|---|---|---|---|---|
| **Method** | **Overclaim** | **Underclaim** | **Overclaim** | **Underclaim** |
| SPECTRA_CC | 5 | 0 | 0 | 0 |
| FASTERCC | 0 | 2 | 0 | 0 |
| FastCC | 0 | 0 | 0 | 0 |
| SwiftCC | 5 | 0 | 20 | 0 |
| Disagreeing reactions: 12 at $10^{-6}$, 20 at $10^{-9}$. | | | | |

Table S2: Dependence of SPECTRA_CC on the solver feasibility tolerance, on AGORA2 community models under a random medium. Values are means over three draws and three timings. Spread is the largest difference in the consistent set across three repeated runs of the *same* model.

| **Community** | **feasTol** | **Runtime (s)** | **LPs** | **Spread** |
|---|---|---|---|---|
| 50 species | $10^{-9}$ | 191 | 10 | 45 |
| | $10^{-8}$ | 2,149 | 155 | 2,350 |
| | $10^{-7}$ | 2,722 | 242 | 2,142 |
| | $10^{-6}$ | 2,918 | 270 | 5,097 |
| 100 species | $10^{-9}$ | 585 | 13 | 129 |
| | $10^{-8}$ | 13,919 | 335 | 19,664 |
| | $10^{-7}$ | 12,547 | 438 | 5,002 |
| | $10^{-6}$ | 13,390 | 476 | 4,150 |

Table S3: Iterated consistency checking on Harvey. Each method was applied to the model, the model was rebuilt from the reactions it reported consistent, and the method applied again. A sound method returns every reaction it is given on the second pass.

| **Method** | **Pass 1** | **Pass 2** | **Pass 3** | **Outcome** |
|---|---|---|---|---|
| SPECTRA_CC | 67,436 (20 LPs) | 0 (2 LPs) | — | collapsed |
| SPECTRA_CC, bounds relaxed | 67,436 (20 LPs) | 67,312 (22 LPs) | 67,312 (22 LPs) | converged |
| FASTERCC | 66,389 (738 LPs) | 66,242 (26 LPs) | 66,242 (24 LPs) | converged |
| Input: 81,094 reactions. Values are consistent reactions returned. | | | | |

Table S4: Summary of the distinct model extraction methods used in constraint-based modeling (CBM) frameworks. CSM: Context-Specific Model, GEM: Genome-Scale Model, MR: Minimal Reactome, MC: Microbial community, MM: Minimal Microbiome.

| **Method** | **CBM-type** | **Constraints** | **Objective** | **Omics data integrated** | **Reference** |
|---|---|---|---|---|---|
| Fastcore | CSM | Stoichiometry; box; Make core reactions consistent | Minimize model size | Transcriptomics | [9] |
| Swiftcore | CSM | Stoichiometry; box; Make core reactions consistent | Minimize model size | Transcriptomics | [8] |
| INIT | CSM | Stoichiometry; box | Maximize inclusion of positively weighted reactions and exclude negatively weighted reactions | Transcriptomics; Proteomics; Metabolomics | [22] |
| MBA | CSM | Stoichiometry; box; Make core reactions consistent | Minimize model size | Transcriptomics | [23] |
| mCADRE | CSM | Stoichiometry; box; Make core reactions consistent | Minimize model size | Transcriptomics | [23] |
| GIMME | CSM | Stoichiometry; box; Flux on biomass reaction | Minimize flux through non-biomass reactions | Transcriptomics | [7] |
| iMAT | CSM | Stoichiometry; box | Maximize inclusion of high confidence reactions and exclude low confidence reactions | Transcriptomics | [24] |
| GIM3E | CSM | Stoichiometry; box; Flux on biomass reaction; bounds on turnover metabolite reactions | Minimize flux through non-biomass reactions | Transcriptomics; Metabolomics | [25] |

| | | | | | |
|---|---|---|---|---|---|
| tINIT | CSM | Stoichiometry; box; Extracted model passes all tasks; Make essential reactions consistent | Maximize inclusion of positively weighted reactions and exclude negatively weighted reactions | Bibliomics; Transcriptomics; Proteomics; Metabolomics | [22] |
| fastGapFill | GEM | Stoichiometry; box; Make core reactions consistent | Minimize model size | Genomics | [26] |
| FastGapFilling | GEM | Stoichiometry; box | Maximize flux through biomass and minimize the flux through the other reactions | Genomics | [27] |
| CarveMe | GEM | Stoichiometry; box; Flux on biomass reaction | Maximize inclusion of positively weighted reactions and exclude negatively weighted reactions | Genomics | [28] |
| gapseq | GEM | Stoichiometry; box | Maximize biomass flux while minimizing flux through other reactions | Genomics | [29] |
| Meneco | GEM | Topology; box; Produce biomass precursors | Minimize model size | Genomics; Transcriptomics; Metabolomics; | [30] |
| MIRAGE | GEM | Stoichiometry; box; Make core reactions and biomass reaction consistent; Growth-associated dilution of all network metabolites | Minimize model size | Genomics | [31] |

| | | | | | |
|---|---|---|---|---|---|
| NetworkReducer | MR | Stoichiometry; box; Make core reactions consistent; Produce the protected metabolites; Extracted model passes all the tasks | Minimize model size | - | [32] |
| minNW | MR | Stoichiometry; box; Make core reactions consistent; Produce the protected metabolites; Extracted model passes all the tasks | Minimize model size | - | [33] |
| Community gap filling | MC | Stoichiometry; box; Flux on biomass reaction | Minimize model size | Genomics | [34] |
| COMMIT | MC | Stoichiometry; box; Flux on biomass reaction | Minimize model size | Genomics | [35] |
| minMicrobiome | MM | Stoichiometry; box; Flux on biomass reaction; Flux on metabolic functionality | Minimize the number of community members | - | [36] |

Table S5: Enriched pathways in Glioblastoma

| S.No | Enriched pathways | Evidence | References |
|---|---|---|---|
| 1 | Eicosanoid metabolism | Pathway-level | [37] |
| 2 | Heme synthesis | Pathway-level | [38] |
| 3 | Starch and sucrose metabolism | Pathway-level | [39] |
| 4 | Sphingolipid metabolism | Pathway-level | [40] |
| 5 | Glycosphingolipid metabolism | Pathway-level | [41] |
| 6 | Fatty acid oxidation | Pathway-level | [42] |
| 7 | Fructose/mannose metabolism | Pathway-level | [43] |
| 8 | Folate metabolism | Pathway-level | [44] |
| 9 | Purine synthesis | Pathway-level | [45] |

Table S6: Enriched pathways in Renal Clear Cell Carcinoma

| S.No | Enriched pathways | Evidence | References |
|---|---|---|---|
| 1 | Chondroitin sulfate degradation | Component-level | [46] |
| 2 | Fatty acid synthesis | Pathway-level | [47] |
| 3 | Glyoxylate and dicarboxylate metabolism | Component-level | [48, 49] |
| 4 | Glycosphingolipid metabolism | Pathway-level | [50] |
| 5 | N-glycan degradation | Pathway-level | [51, 52] |
| 6 | Hyaluronan metabolism | Pathway-level | [53, 54] |
| 7 | Starch and sucrose metabolism | Component-level | [55] |

Table S7: Enriched pathways in Liver cancer

| S.No | Enriched pathways | Evidence | References |
|---|---|---|---|
| 1 | Keratan sulfate degradation | Component-level | [56] |
| 2 | Fatty acid synthesis | Pathway-level | [57] |
| 3 | Heme synthesis | Pathway-level | [58] |
| 4 | Fatty acid oxidation | Pathway-level | [59] |
| 5 | Chondroitin sulfate degradation | Component-level | [56] |
| 6 | Glycosphingolipid metabolism | Component-level | [60] |
| 7 | Starch and sucrose metabolism | Pathway-level | [61] |
| 9 | Sphingolipid metabolism | Pathway-level | [62] |
| 11 | Glyoxylate and dicarboxylate metabolism | Component-level | [63] |
| 12 | Drug metabolism | Pathway-level | [64] |
| 13 | Urea cycle | Pathway-level | [65] |
| 14 | Glycerophospholipid metabolism | Pathway-level | [66] |
| 15 | Miscellaneous | Component-level | [67, 68] |
| 16 | N-glycan degradation | Pathway-level | [69] |
| 16 | Eicosanoid metabolism | Pathway-level | [70] |
| 17 | Phenylalanine metabolism | Pathway-level | [63, 71] |
| 18 | Nucleotide interconversion | Pathway-level | [72, 73] |
| 19 | Steroid metabolism | Pathway-level | [74] |

Table S8: Enriched pathways in Lung Adenocarcinoma

| S.No | Enriched pathways | Evidence | References |
|---|---|---|---|
| 1 | Heme synthesis | Pathway-level | [75] |
| 2 | Fatty acid oxidation | Pathway-level | [76, 77] |
| 3 | Lysine metabolism | Component-level | [78] |
| 4 | Drug metabolism | Pathway-level | [79] |
| 5 | Citric acid cycle | Pathway-level | [80] |
| 6 | Glycosphingolipid metabolism | Pathway-level | [81] |
| 7 | Heme degradation | Pathway-level | [75] |
| 8 | Cholesterol metabolism | Pathway-level | [82, 83] |

Table S9: Enriched pathways in Retinoblastoma

| S.No | Enriched pathways | Evidence | References |
| --- | --- | --- | --- |
| 1 | Keratan sulfate synthesis/degradation | Component-level | [84] |
| 2 | Chondroitin synthesis/Chondroitin sulfate degradation | Component-level | [84] |
| 3 | Blood group synthesis | Uncertain | - |
| 4 | Sphingolipid metabolism | Pathway-level | [85, 86] |
| 5 | Fatty acid oxidation | Pathway-level | [87] |
| 6 | Eicosanoid metabolism | Component-level | [88] |
| 7 | Heme synthesis | Pathway-level | [89] |
| 8 | Cholesterol metabolism | Pathway-level | [90] |
| 9 | Glycosphingolipid metabolism | Pathway-level | [91, 86] |
| 10 | O-glycan metabolism | Component-level | [92, 91] |
| 11 | Hyaluronan metabolism | Pathway-level | [93, 94] |
| 12 | Bile acid synthesis | Component-level | [85, 88] |

Table S10: Enriched pathways in Pancreatic Adenosquamous Carcinoma

| S.No | Enriched pathways | Evidence | References |
| --- | --- | --- | --- |
| 1 | Fatty acid oxidation | Pathway-level | [95] |
| 2 | Eicosanoid metabolism | Pathway-level | [96] |
| 3 | Fatty acid synthesis | Pathway-level | [97] |
| 4 | Cholesterol metabolism | Pathway-level | [98] |
| 5 | Glycosphingolipid metabolism | Pathway-level | [99] |

Table S11: Enriched pathways in Prostate Small Cell Carcinoma

| S.No | Enriched pathways | Evidence | References |
|---|---|---|---|
| 1 | Fatty acid synthesis | Pathway-level | [100] |
| 2 | Fatty acid oxidation | Pathway-level | [101] |
| 3 | Drug metabolism | Component-level | [102] |
| 4 | Glycosphingolipid metabolism | Pathway-level | [103, 104] |
| 5 | Starch and sucrose metabolism | Component-level | [105] |
| 6 | Glyoxylate and dicarboxylate metabolism | Pathway-level | [106] |
| 7 | CoA synthesis | Uncertain | - |

Table S12: Enriched pathways in Invasive Breast Carcinoma

| S.No | Enriched pathways | Evidence | References |
|---|---|---|---|
| 1 | Eicosanoid metabolism | Pathway-level | [107] |
| 2 | Purine synthesis | Pathway-level | [108] |
| 3 | Fatty acid oxidation | Pathway-level | [109] |
| 4 | Bile acid synthesis | Pathway-level | [106] |
| 5 | O-glycan metabolism | Pathway-level | [110] |
| 6 | Drug metabolism | Pathway-level | [111] |
| 7 | Vitamin D metabolism | Pathway-level | [112] |
| 8 | Citric acid cycle | Pathway-level | [113] |
| 9 | Cholesterol metabolism | Pathway-level | [114] |

Table S13: Enriched pathways in High-Grade Serous Ovarian Cancer

| S.No | Enriched pathways | Evidence | References |
|---|---|---|---|
| 1 | Eicosanoid metabolism | Pathway-level | [115] |
| 2 | Fatty acid oxidation | Pathway-level | [116] |
| 3 | Steroid metabolism | Pathway-level | [117] |
| 4 | Glutathione metabolism | Pathway-level | [118] |
| 5 | Drug metabolism | Pathway-level | [119] |
| 6 | Cholesterol metabolism | Pathway-level | [120] |

Table S14: Enriched pathways in Gallbladder Cancer

| **S.No** | **Enriched pathways** | **Evidence** | **References** |
|---|---|---|---|
| 1 | Keratan sulfate synthesis / degradation | Uncertain | - |
| 2 | Chondroitin synthesis / Chondroitin-sulfate degradation | Uncertain | - |
| 3 | Blood group synthesis | Component-level | [121, 122] |
| 4 | Heparan sulfate degradation | Pathway-level | [123, 124] |
| 5 | Sphingolipid metabolism | Pathway-level | [125] |
| 6 | Heme synthesis | Uncertain | - |
| 7 | O-glycan metabolism | Component-level | [126] |
| 8 | Fatty acid oxidation | Pathway-level | [127, 128] |
| 9 | Starch and sucrose metabolism | Uncertain | - |
| 10 | Glycosphingolipid metabolism | Component-level | [129, 130] |
| 11 | Fatty acid synthesis | Pathway-level | [131, 132] |
| 12 | Glyoxylate and dicarboxylate metabolism | Component-level | [133] |
| 13 | Cholesterol metabolism | Pathway-level | [134] |

### S3 Supporting Figures

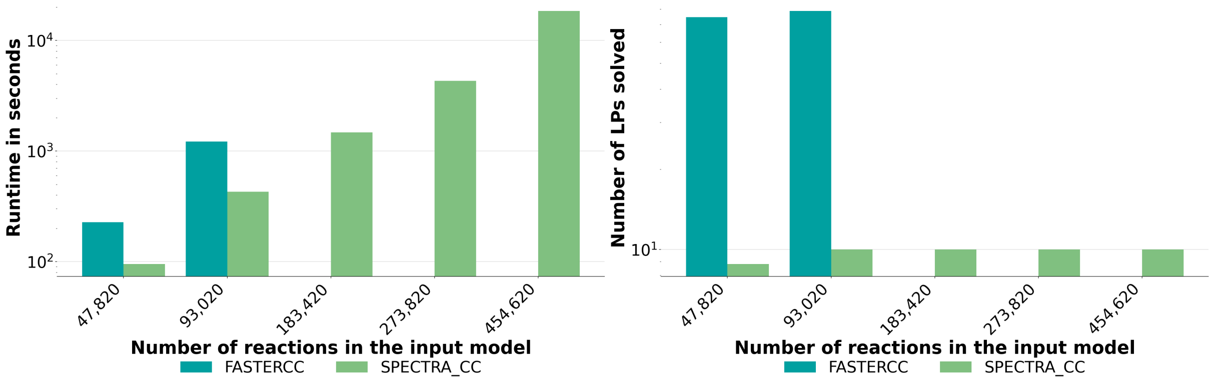

Figure S8: **SPECTRA_CC solves a constant number of LPs as model size grows.** Expanded Recon3D reconstructions from 47,820 to 454,620 reactions, with 500-10,000 reactions deleted at random; bars are means over three draws and three timings. SPECTRA_CC requires 10 LPs at every size from 93,020 reactions upward. FASTERCC has no bar beyond 93,020 reactions because it returns no result at those sizes.

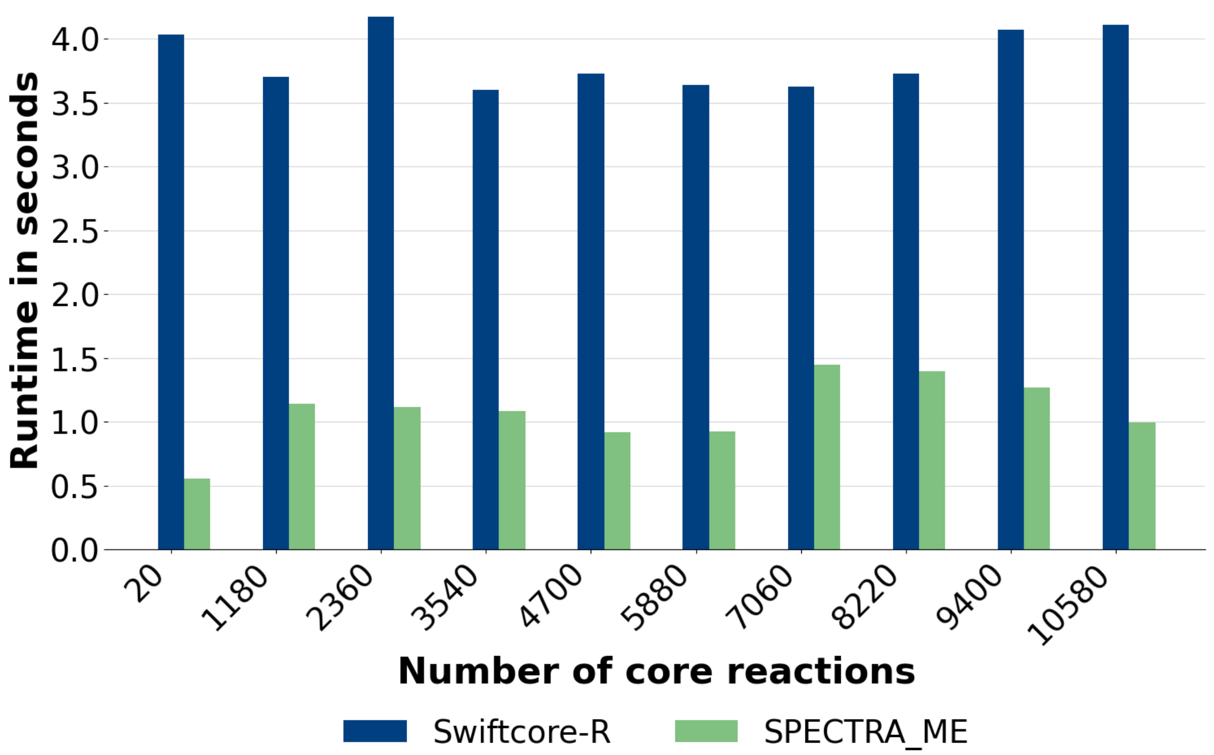

Figure S9: SPECTRA_ME is the fastest algorithm when compared to Swiftcore's implementation with reduction

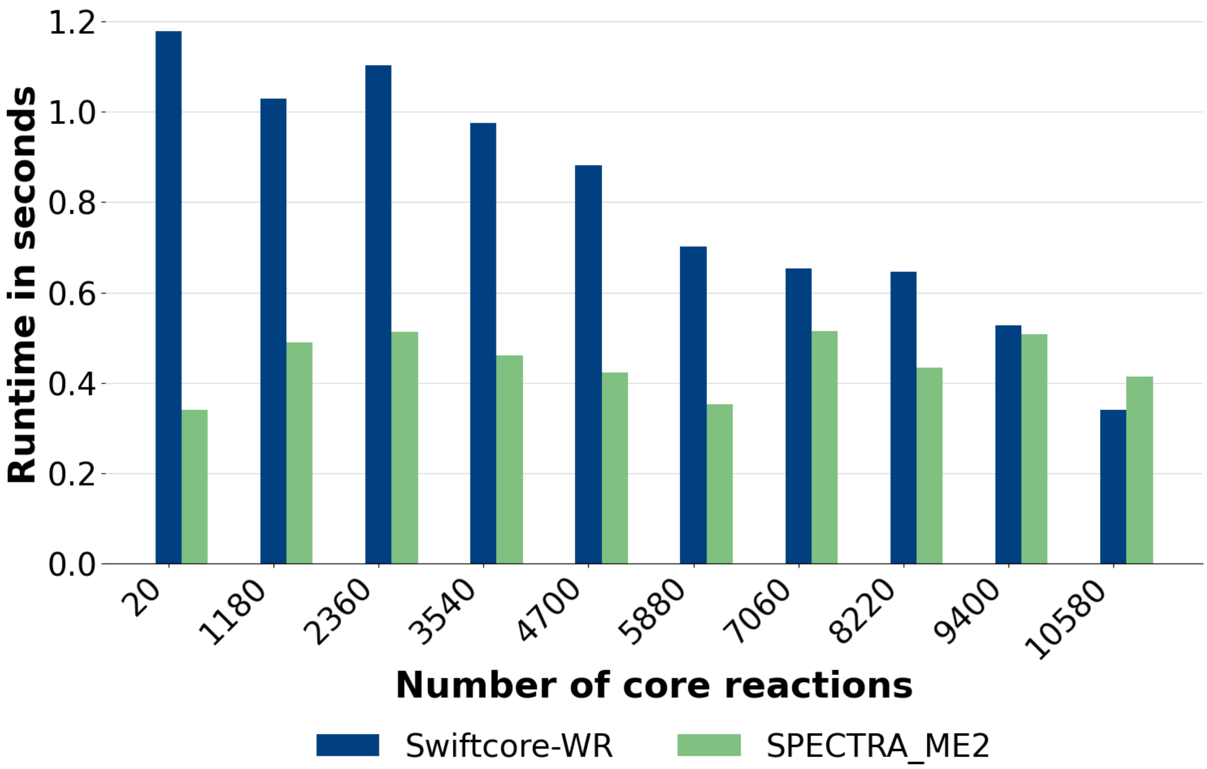

Figure S10: **SPECTRA_ME2 is efficient algorithm compared to Swiftcore without reduction (WR) (a)** SPECTRA_ME2 accelerates the generation of metabolic models when the constraints on non-core reversible reactions are not considered. SPECTRA_ME2 is similar to SPECTRA_ME except non-consideration of the constraints on non-core reversible reactions.

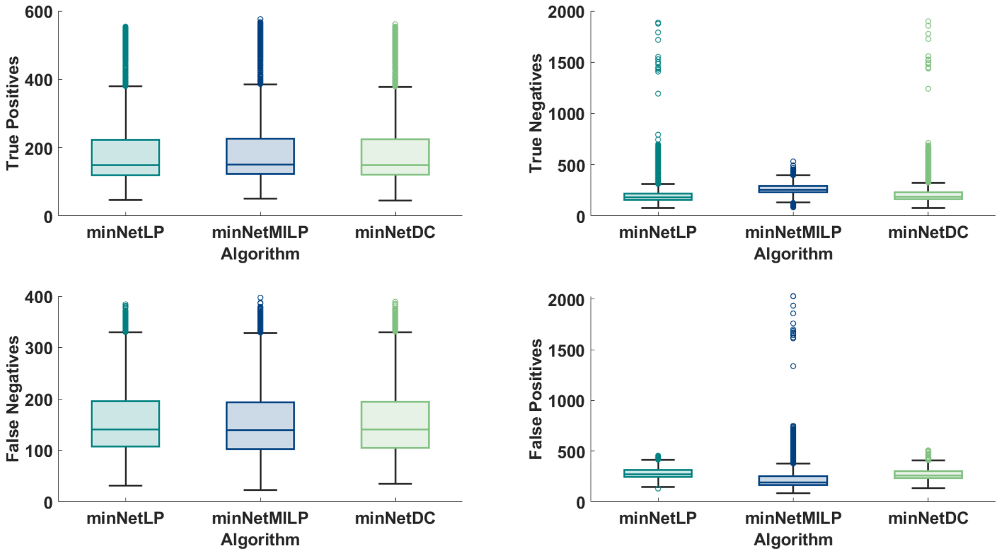

Figure S11: **Comparison of minimal-network–based inference methods:** All algorithms show similar performance for true positives and false negatives. For true negatives and false positives, minNetMILP outperforms the other methods. Statistical significance was confirmed using a right-tailed Wilcoxon signed-rank test for true negatives ($p = 0$) and a left-tailed Wilcoxon signed-rank test for false positives ($p = 0$).

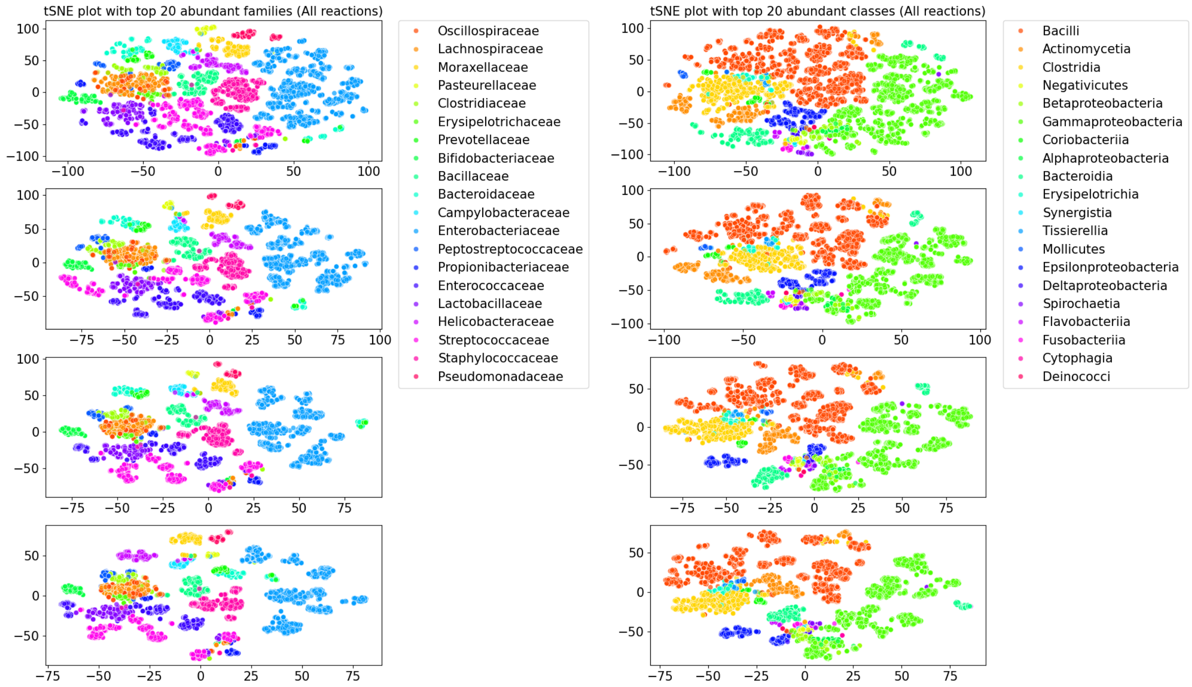

Figure S12: tSNE plots for the gap-filled AGORA2 models with varying perplexity values. The distinct colors refer to the top-20 abundant families (left-panel) or top-20 abundant classes (right-panel). Here all the reactions in the final gap-filled AGORA2 model is taken into account.

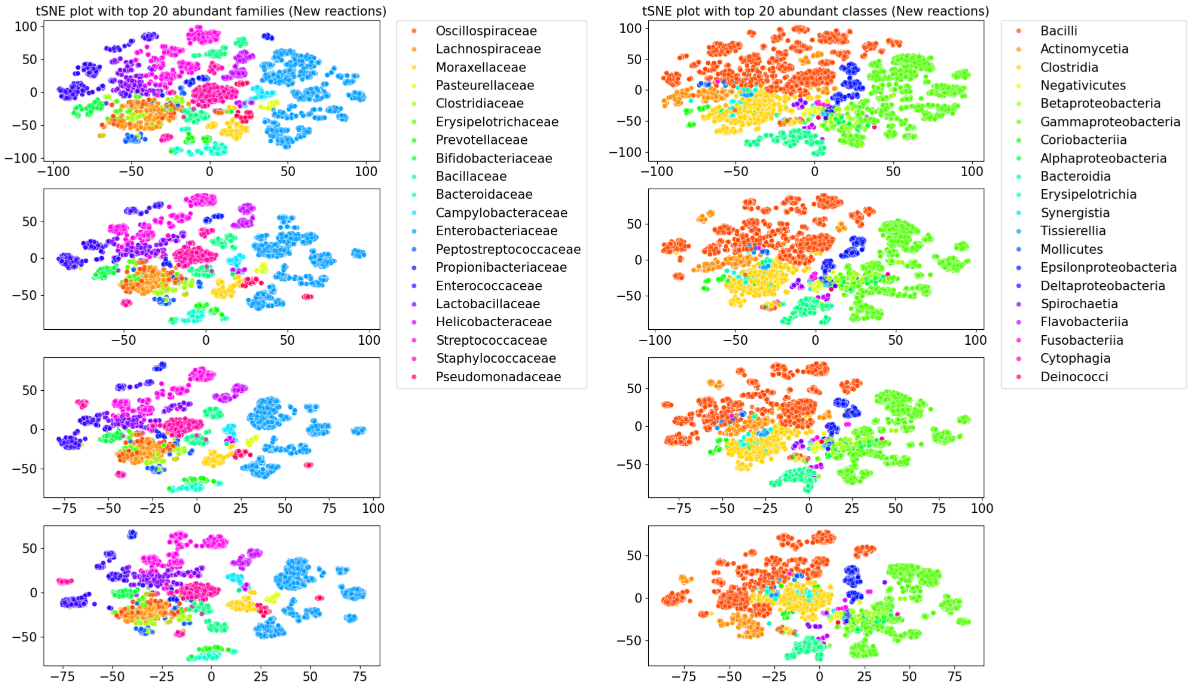

Figure S13: tSNE plots for the gap-filled AGORA2 models with varying perplexity values. The distinct colors refer to the top-20 abundant families (left-panel) or top-20 abundant classes (right-panel). Here only the newly added reactions in the final gap-filled AGORA2 model is taken into account.

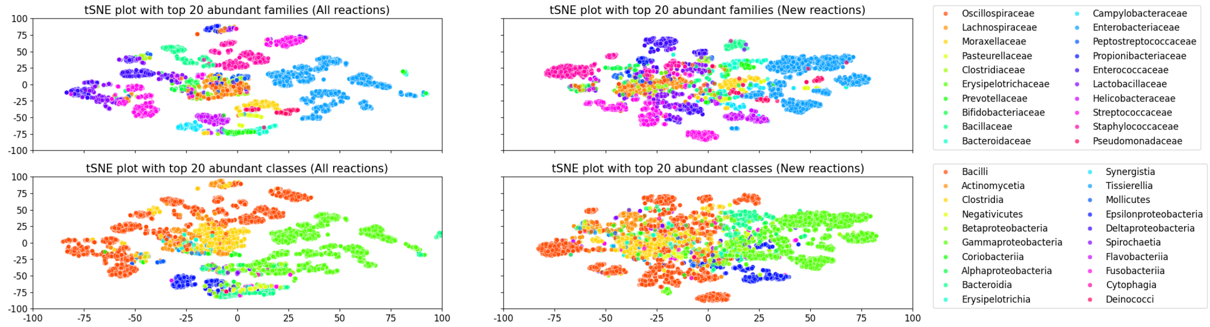

Figure S14: tSNE clustering of the subsystems in newly built models and the newly added reactions, reveal the taxonomic patterns underlying these models.
